# Distinct pathways of adaptive evolution in *Cryptococcus neoformans* reveal a point mutation in adenylate cyclase with drastic tradeoffs for pathogenicity

**DOI:** 10.1101/2022.09.27.509772

**Authors:** Zoë A. Hilbert, Krystal Y. Chung, Joseph M. Bednarek, Mara W. Schwiesow, Jessica C.S. Brown, Nels C. Elde

## Abstract

Pathogenic fungi populate a wide range of environments and infect a diversity of host species. Despite this substantial biological flexibility, the impact of interactions between fungi and their hosts on the evolution of pathogenicity remains unclear. We studied how repeated interactions between the fungus *Cryptococcus neoformans* and relevant environmental and mammalian host cells—amoeba and mouse macrophages—shape the evolution of this model fungal pathogen. First, using a collection of clinical and environmental isolates of *C. neoformans*, we characterized a range of survival phenotypes for these strains when exposed to host cells of different species. We then performed serial passages of an environmentally isolated *C. neoformans* strain through either amoeba or macrophages for ~75 generations to observe how these interactions select for improved replication within hosts. In an adapted population, we identified a single point mutation in the adenylate cyclase gene, *CAC1*, that swept to fixation and confers a strong competitive advantage for growth inside of macrophages. Strikingly, this growth advantage in macrophages is inversely correlated with disease severity during mouse infections, suggesting that adaptations to specific host niches can markedly reduce the pathogenicity of these fungi. These results raise intriguing questions about the influence of cAMP signaling on pathogenicity and highlight the role of seemingly small adaptive changes in promoting fundamental shifts in the intracellular behavior and virulence of these important human pathogens.

## Introduction

The ability to adapt and thrive in different environments is critical for the evolutionary success of most organisms. For pathogenic microbes, adaptive traits that facilitate survival within host cells or tissues primarily evolve through repeated interactions with host immune factors or exposure to host-like conditions. However, for environmentally-derived pathogens, such as most fungi, the mechanisms by which these organisms initially gain the ability to infect mammalian hosts are not well understood. Furthermore, our understanding of how fungi, in particular, might adapt over short timescales within animal hosts in response to pressures from host immune cells, such as macrophages, is similarly limited.

The human fungal pathogen *Cryptococcus neoformans*, is a globally distributed haploid yeast which is ubiquitous in the environment and often found associated with trees or bird guano^1^. Human infection with *C. neoformans* occurs primarily through the inhalation of fungal spores or dessicated yeasts into the lungs where, in immune competent patients, infection is either cleared or contained in a persistent state for long periods of time^2^. Symptomatic disease is observed mostly in immune compromised patients, where dissemination to the brain leads to cryptococcal meningitis, which is lethal unless treated and is currently estimated to be responsible for more than 115,000 deaths annually^3^. Though *C. neoformans* is an effective and important human pathogen, identifying the evolutionary pressures and genomic adaptations that led to the emergence of pathogenicity in this species has remained elusive.

Some evolutionary studies of virulence in *C. neoformans* have focused on characterization of large collections of isolates obtained from both clinical and environmental sampling^4–10^. By combining whole genome sequencing of these strains with phenotypic profiling of pathogenicity-associated traits such as melanization, growth at high temperature, or success in *in vivo* infection models, genome-wide association studies have identified genomic changes that may underlie the increased success of some of these strains as pathogens^4,9,10^. While such studies are critical in identifying key genes and pathways associated with increased pathogenicity, they are less useful in identifying the selective pressures that drove the acquisition of these traits. Instead, complementary studies have focused on the role of environmental hosts in shaping the evolution of virulence in *Cryptococcus* species. This intriguing body of work rests on a hypothesis that posits that the evolution of virulence in environmental fungi can be explained by similarities in selective pressures imposed by environmental predators, such as amoebae and nematodes, and those experienced during animal infections, including in humans^11,12^.

Amoebae have been particularly well studied as a potential source of environmental selective pressure on diverse microbes given their similarities to the macrophages of animal immune systems. Like macrophages, amoebae can phagocytose microorganisms, and undergo many similar processes following phagocytosis including phagosome maturation and acidification^13^. Studies of bacterial species, like *Legionella pneumophila,* have identified virulence factors that are required for success in both specific amoeba species and mammalian host cells, providing strong evidence that similarities across amoeba and mammalian host cell environments could select for broadly effective virulence strategies^14–17^. For *C. neoformans*, capsule production has been shown to have protective roles against both amoeba and mammalian hosts, and numerous other features of the intracellular behavior of *C. neoformans* appear to be conserved across these evolutionarily diverse species^18–21^. However, whether pathogenicity can truly be evolved through repeated interactions with amoebae or other environmental hosts and the mechanisms through which this occurs have not been well studied.

Beyond environmental interactions, adaptation within mammalian hosts also has the potential to select for genetic variation within fungal populations with important implications for disease progression. “Microevolution” studies performed by analyzing serial samples from patients with recurrent cryptococcal meningitis has revealed numerous adaptive phenotypic changes in these strains over the course of months of exposure to the host environment; though correlation between genetic changes and these adaptive phenotypes has proven challenging to assess beyond identifying the mechanisms of resistance to antifungal treatments^22–24^. Serial *in vitro* passaging schemes, on the other hand, such as those performed with the fungal pathogen *Candida glabrata*, suggest this to be a powerful system to identify phenotypic changes involved in host adaptation *and* identify the molecular mechanisms associated with such changes^25–28^. Few such studies have compared the adaptive changes that occur in response to exposure to different host species. In addition, most experiments have focused on changes that occur in commonly used laboratory strains or closely related clinical isolates, which may not accurately capture the evolutionary trajectories taken by divergent isolates of these fungal species.

Here, we used clinical and environmental isolates of *C. neoformans* to compare the survival of strains from different genetic backgrounds following phagocytosis by either environmental (amoeba) or mammalian (mouse macrophage) hosts. We then serially passaged one of these environmental isolates through amoeba or macrophages to isolate host-adapted strains with enhanced abilities to survive in the intracellular niches of highly diverged host organisms. Remarkably, through these approaches, we discovered that only a single missense mutation in the *CAC1* adenylate cyclase gene is sufficient to confer enhanced macrophage replication and can modulate the balance between host adaptation and pathogenicity in this important human fungal pathogen.

## Results

### Natural variation in C. neoformans strain replication in amoebae and macrophages

We profiled the abilities of 14 unique *C. neoformans* isolates from sub-Saharan Africa to replicate either in cells of the amoeba species, *Acanthamoeba castellanii,* or in the J774A.1 mouse macrophage-like cell line (Figure 1). The strains chosen represent three of the four major non-recombining *C. neoformans* sub-lineages—VNI, VNBI, and VNBII—and are predominantly environmental isolates, though several clinical isolates were also included for comparison (Figure 1A)^4^.

**Figure 1.**
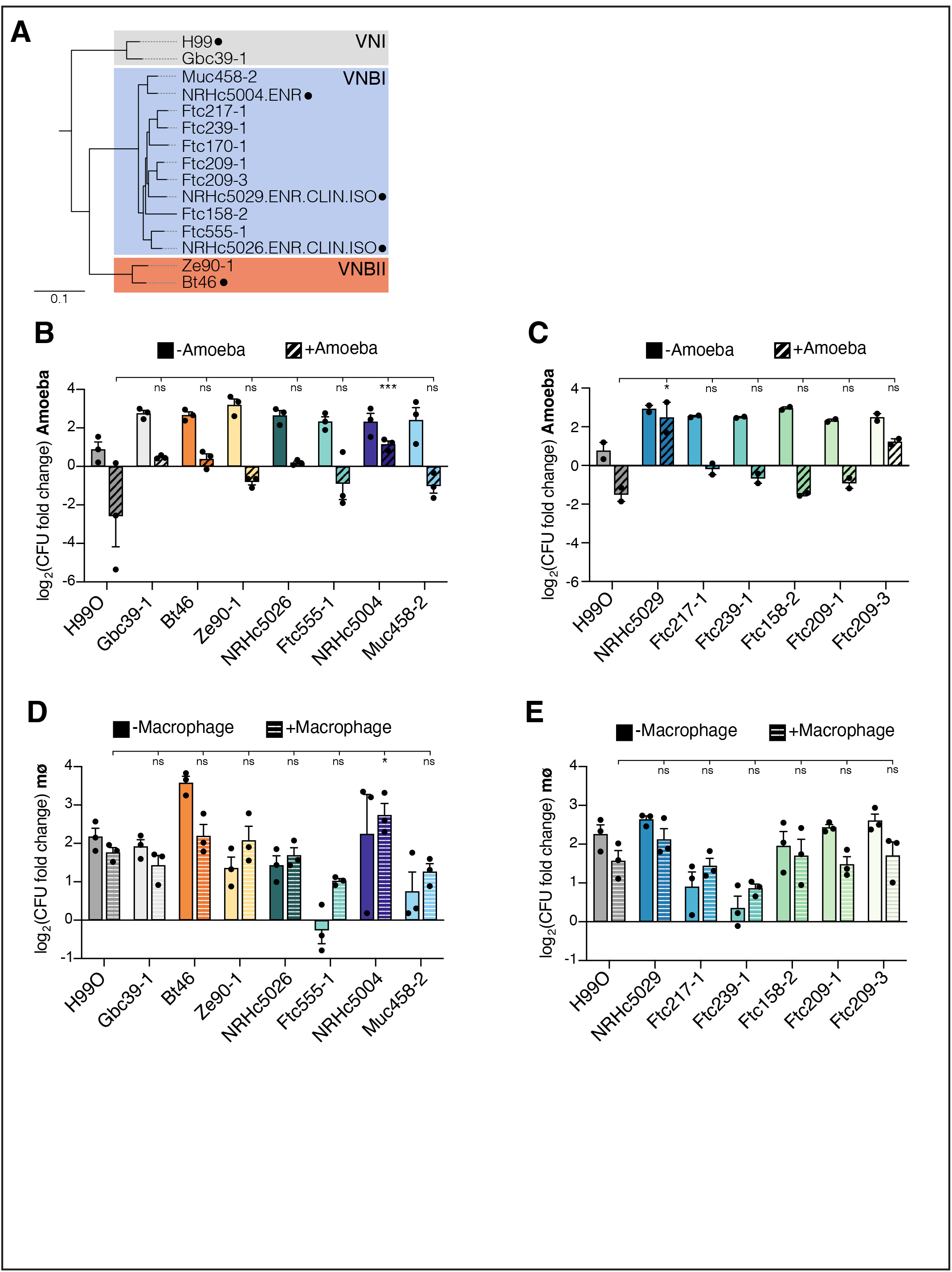
Natural variation of *C. neoformans* clinical and environmental isolates survival in amoebae or macrophage hosts. (A) Phylogenetic tree of clinical and environmental *C. neoformans* strains used in this paper. Tree was extracted from a larger published phylogeny using the Treehouse R-package^4,67^. Lineages are indicated by colored boxes and labels, and black dots indicate clinical isolates. All other strains were isolated from environmental sources. Scale bar indicates 0.1 substitutions/site. (B) and (C) Replication of *C. neoformans* strains in co-incubation experiments with *A. castellanii* amoeba cells. Solid bars indicate the growth of the strain in minimal Ac Medium without amoeba cells present. Striped bars indicate strain growth with amoeba present. (D) and (E) Replication of the same strains in co-incubation experiments with J774A.1 macrophage cells. Solid bars indicate growth of the strains in macrophage growth medium without macrophage cells. Striped bars indicate replication with macrophages present. For (B)-(E), bars show the average values ± SEM across 2-3 independent experiments on different days. Dots indicate the average values for 3 replicates for each independent experiment. Significance was assessed by comparison to the H99O reference strain by ordinary one-way ANOVA followed by Dunnett’s multiple comparisons test. * p<0.05, *** p<0.0001, ns not significant.

When these strains were incubated with *A. castellanii* cells for 24 hours, we observed robust killing of the cryptococcal cells in nearly all of the strains tested, despite roughly similar growth in the absence of amoebae (Figure 1B and C). Notably, we identified two strains with significantly enhanced abilities to replicate inside of the amoeba cells—NRHc5004 and NRHc5029—both of which were isolated from clinical sources. In contrast to the killing we observed in co-culture experiments with *A. castellanii*, all of the tested *C. neoformans* isolates were able to replicate to some extent when incubated with mouse macrophages (Figure 1D and E). The clinical isolate NRHc5004 again had an enhanced ability to replicate in macrophages, underscoring the possibility that there may be some strategies for intracellular replication that confer success in the evolutionarily diverse host cells tested here.

One explanation for strain-level variation in host cell survival and replication could be that there are differences in the phagocytosis of these *C. neoformans* strains by host cells. To account for this possibility, we determined the percent phagocytosis for each of the strains by comparing the amount of the input culture remaining after an initial phagocytosis step in our co-incubation experiments (Supplemental Figure 1). We observe no significant differences in phagocytosis across these strains in either the amoeba or macrophage experiments. However, we do note that phagocytosis was increased in the macrophage experiments overall (>80% of cells phagocytosed), likely due to the fact that these cells are opsonized with anti-capsule antibody prior to their addition to macrophage cultures (Supplemental Figure 1C and D). Regardless, differences in phagocytosis are not contributing significantly to the observed intracellular replication phenotypes.

In addition to strain-specific variation in macrophage replication, there was even more pronounced variation in the ability of these strains to replicate under tissue culture conditions in the absence of macrophages (solid bars, Figure 1D and E). In particular, we observed that the environmental isolate Ftc555-1 was killed by incubation in tissue culture media alone, but that this could be partially rescued through the addition of macrophage cells to the culture (Figure 1D). The low levels of media replication that we observed in Ftc555-1 highlighted this strain as an ideal candidate for serial passaging and host adaptation studies. We reasoned that we could exploit the preference of this strain for the macrophage intracellular environment to ask questions about how the evolution of this strain is shaped by host selective pressures in the absence of excessive growth outside of these cells.

### Serial passaging of C. neoformans through host cells selects for strains with enhanced survival

To assess the ability of *C. neoformans* to adapt to the intracellular environments of diverse host cells, we devised a serial passaging scheme where Ftc555-1 was repeatedly subjected to phagocytosis by either *A. castellanii* amoebae or J774A.1 macrophages. This was followed by a 24 hour-long incubation during which the fungi could replicate inside of the host cell and an outgrowth step to recover enough yeast cells to set up the next round of passaging (Figure 2A). The experiment was performed in parallel using both amoebae and macrophage host cells; three independent populations were evolved in each cell type and are hereafter referred to as lines M1-M3 (for macrophage-passaged) and A1-A3 (for amoeba-passaged). These replicate populations were treated almost identically, with one critical difference: in two of the three populations (lines M1/M2 and A1/A2), all extracellular yeast were discarded after the 24 hour co-incubation step to enrich for the intracellular and host cell-associated population, while in the third line (M3 and A3) all yeast cells were collected, including those that had escaped from the host cell into the media (see Experimental Methods).

**Figure 2.**
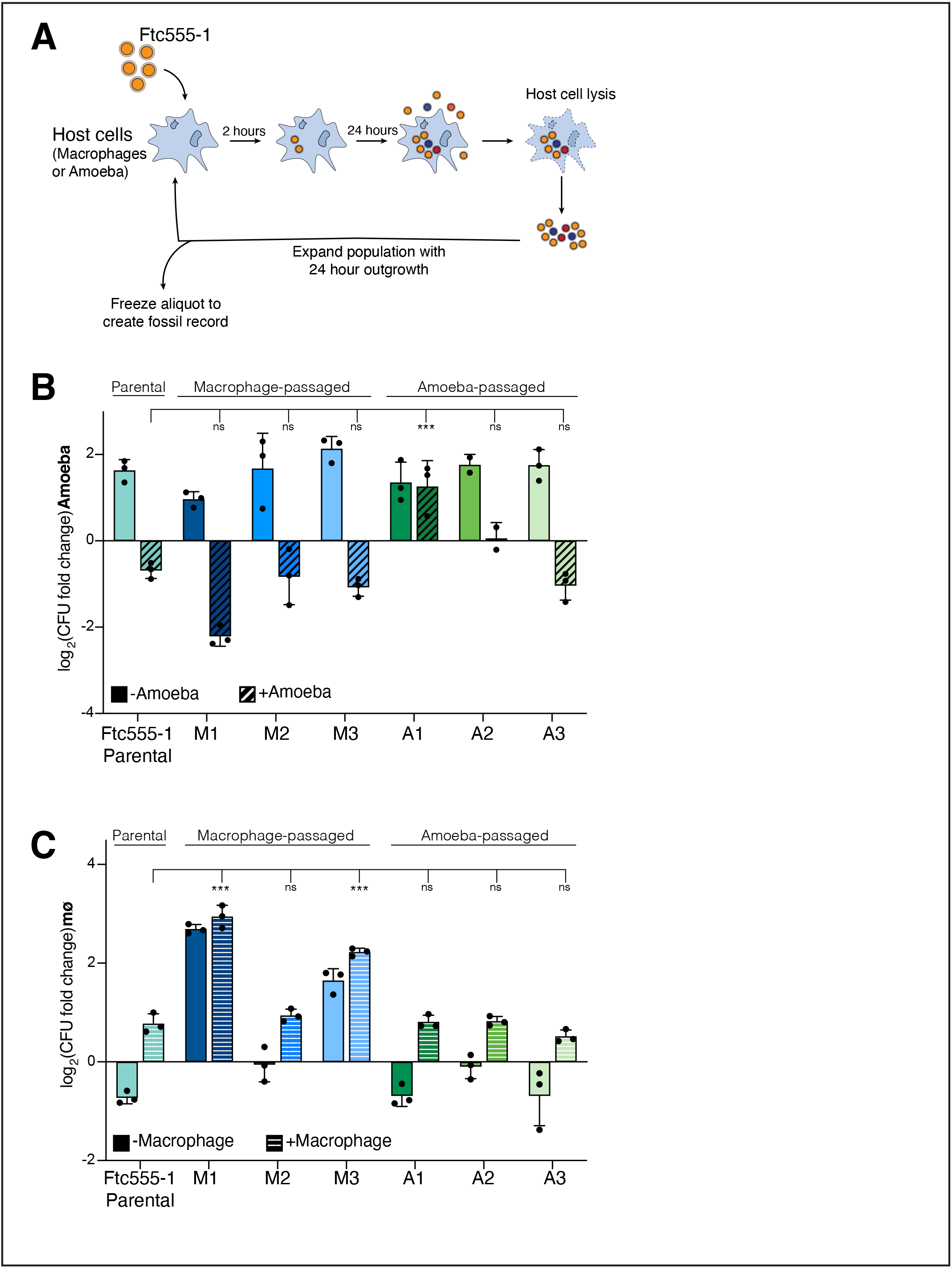
Serial passaging of *C. neoformans* through macrophages or amoeba cells selects for host adapted strains. (A) Schematic of the serial passaging set up used in this study. (B) Replication phenotypes of passaged strains in co-culture with amoeba cells. Single colonies from each individually evolved population were tested for growth in amoeba. Strains M1-M3 were passaged through macrophages, A1-A3 were passaged through amoeba. Solid bars indicates media only growth; striped bars indicate growth following amoeba phagocytosis. (C) Replication phenotypes of passaged strains in co-culture with mouse macrophage-like cells. The same single colonies were tested as in (B) and data is plotted identically. For (B) and (C), plotted data is from one representative experiment. Error bars indicate SD and each dot indicates the value for one of the three biological replicates from that experiment. Significance was assessed between the parental strain values and all evolved strains by ordinary one-way ANOVA followed by Dunnett’s multiple comparisons test. *** p<0.0001, ns not significant.

We ran courses of experimental evolution for a total of 12 passages over the span of roughly one month of continuous culturing. At the end of the experiment, we collected the full population and froze down pooled cultures. Importantly, aliquots of each passage were frozen over the course of the entire experiment to create a “fossil record,” which could be thawed, as necessary, to track the acquisition of adaptive mutations over time and assess intermediate phenotypes.

Following the final passage, we analyzed the ability of individual colonies from each population to replicate when cultured with either amoebae or macrophage hosts (Figure 2B and C). When assessed in culture with amoebae, only a single evolved population showed significant changes in replication: the amoeba-passaged A1 line (Figure 2B). The enhanced replication of this strain is specific to growth in culture with host cells, as the media-only replication rates remained unchanged from the parental strain. In macrophages, we observed enhanced replication in two of the three macrophage-passaged isolates, with virtually no change in phenotype for any of the other evolved strains (Figure 2C). Notably, in both the strains with enhanced replication in macrophages (M1 and M3), we also observed a marked improvement in growth under tissue culture conditions in the absence of macrophages (Figure 2C, solid bars), a phenotype that was not observed for any of the other evolved strains across both host cell types. Importantly, we assessed phagocytosis of these evolved strains by both amoebae and macrophages and observed that there were minimal differences compared to the parental strain, indicating that the phenotypes we observe are independent of changes in phagocytosis (Supplemental Figure 2).

### A macrophage-evolved strain outcompetes the parental for growth inside of macrophages

Of the strains recovered from our serial passaging approach, the M1 strain stood out as having a robust host-adapted phenotype, with a nearly 5-fold increase in replication in macrophages compared to the parental strain (Figure 2C). We reasoned that a more detailed analysis of the adaptive phenotype of this M1 strain and the mechanism by which it arose was warranted. Such analysis would have important implications for our understanding of the types of changes that are selected for by macrophage interactions.

Our initial characterizations of the M1 strain were performed on isogenic cultures of this strain, but mutations that contribute to adaptive phenotypes must confer a fitness advantage within a population of heterogeneous cells. We sought to determine the relative fitness advantage of this M1 isolate when compared to the parental Ftc555-1 by directly competing these strains against each other in our macrophage experiments. To do this, we generated a version of the parental strain labeled with a nourseothricin-resistance cassette (NAT^R^) which we competed against unlabeled strains of interest in co-incubation experiments with macrophages. Differential plating of the recovered populations from these experiments onto plates with or without nourseothricin allowed us to determine the relative growth advantage of each strain.

When the M1 strain was competed against the NAT^R^ parental strain, we observed that the evolved M1 strain was able to outcompete the parental strain by over 5-fold in macrophages and nearly 40-fold in the media-only conditions (Figure 3A). In contrast, when the parental Ftc555-1 was instead competed against the NAT^R^-labeled version of itself as an important control, we observed a competitive index of ~1 under all conditions (Figure 3A). Therefore, the evolved M1 isolate has a strong fitness advantage over parental Ftc555-1 and this advantage applies to both growth in the macrophage as well as under tissue-culture conditions without host cells present.

**Figure 3.**
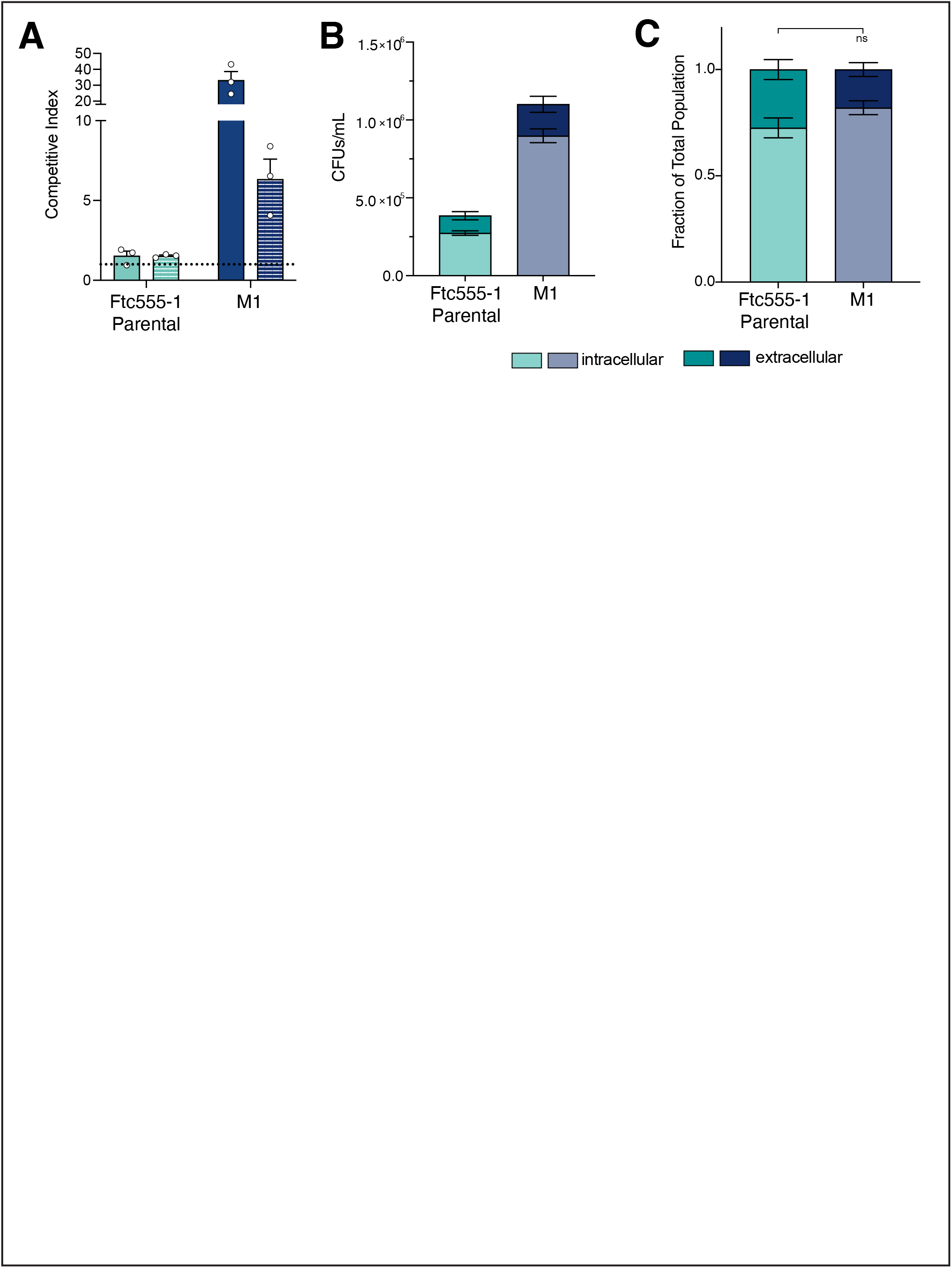
The M1 evolved strain outcompetes the Ftc555-1 parental strain in macrophages. (A) Competition experiments between the NAT^R^-labeled parental strain and unlabeled parental (teal), and M1 (navy blue) strains. Competitive indices are calculated as the ratio of the output samples divided by the ratio of the input samples (see Experimental Methods), with a CI=1 (dotted line) indicating equal growth of the two strains. Solid bars indicate competitive indices in media only, solid bars indicate competitive indices in the macrophage. Plotted values indicate the average value ± SEM of three replicate experiments carried out on three separate days; dots indicate the average value of replicates from a single experiment. (B) Total CFU counts of Ftc555-1 and M1 strains following macrophage co-incubation. Intracellular (light shades) and extracellular (dark shades) populations were collected and plated separately from experiments. Bars show the average values ± SEM from two independent experiments on separate days. (C) Intracellular and extracellular growth following macrophage co-incubation plotted as a fraction of the total population. The same data from (B) but presented as a fraction rather than as raw CFU counts. Significance was assessed by unpaired t-test with Welch’s correction.

The strength of the competitive advantage we observe for M1 in the media-only conditions raises the possibility that the increased macrophage replication of this strain may simply reflect enhanced extracellular growth following escape from macrophages, either through killing of the host cell or non-lytic exocytosis^29–31^. To address this possibility, we assessed the role of extracellular growth in contributing to the M1 enhanced growth phenotype through several methods. First, we determined the amount of extracellular growth that occurs in the parental and M1 evolved strains under our normal experimental conditions. We performed co-incubation experiments with macrophages where, instead of harvesting all fungal cells collectively, we separated intracellular and extracellular populations and enumerated the CFUs from each sub-population. The M1 CFUs from these experiments were significantly higher than the ancestral strain for both the intra- and extracellular populations (Figure 3B). However, when assessed as a fraction of the total recovered CFUs, there was no significant difference in the percentage of the population that was extracellular at the end of the experiment (Figure 3C). This demonstrates that the M1 strain is not escaping macrophages at a noticeably higher rate than the Ftc555-1 parent, and extracellular replication is not a confounding issue at this time point.

We also assessed the possible contribution of extracellular growth to our evolved phenotype in the M1 strain by using the antifungal drug fluconazole to limit *C. neoformans* replication in the media (Supplemental Figure 3). Following phagocytosis by macrophages, *Cryptococcus* cells are somewhat protected from the effects of fluconazole, which allowed us to perform competition experiments between M1 and parental strains where replication was limited to the macrophage intracellular environment (Supplemental Figure 3B and C).

Competition experiments with fluconazole demonstrate that the fitness advantage we observe in M1 is due to an enhanced ability to replicate in the intracellular niche; the M1 strain had a nearly identical competitive index (>5-fold advantage) compared to the parental strain when grown in the presence *or* absence of fluconazole (Figure 3C and Supplemental Figure 3C). Despite the fact that we observed no difference in the minimum inhibitory concentration (MIC) for fluconazole between the two strains, the evolved M1 strain also maintained its fitness advantage in media with fluconazole present, though at a much reduced level (Supplemental Figure 3). Combined, these data demonstrate that our experimental evolution selected for a strong intracellular fitness advantage in the M1 macrophage-passaged strain.

### *Enhanced growth of the M1 strain is caused by a single nucleotide polymorphism in the* CAC1 *adenylate cylase gene*

To identify the adaptive genetic changes underlying the phenotype of the M1 strain, we used a combination of Nanopore and Illumina sequencing to assemble a high-quality reference genome of the Ftc555-1 starting isolate. We then performed whole genome sequencing on several individual colonies from the M1 population as well as on a pooled culture started from the frozen glycerol stock of the final passage from our experimental evolution. Surprisingly, we found only a single nucleotide polymorphism that was shared across all these samples that differentiated the M1 strain from the Ftc555-1 parental isolate. This mutation, which was fixed in both the single colonies as well as the pooled culture, changes a single amino acid— Arg1227Pro—in the gene *CAC1* (CNAG_03202). *CAC1* encodes the adenylate cyclase gene in *C. neoformans*, which is required for the production of cyclic adenosine monophosphate (cAMP) and the regulation of numerous aspects of *Cryptococcus* biology, including mating, cell size, and capsule and melanin production, among others (Figure 4A)^32–36^.

**Figure 4.**
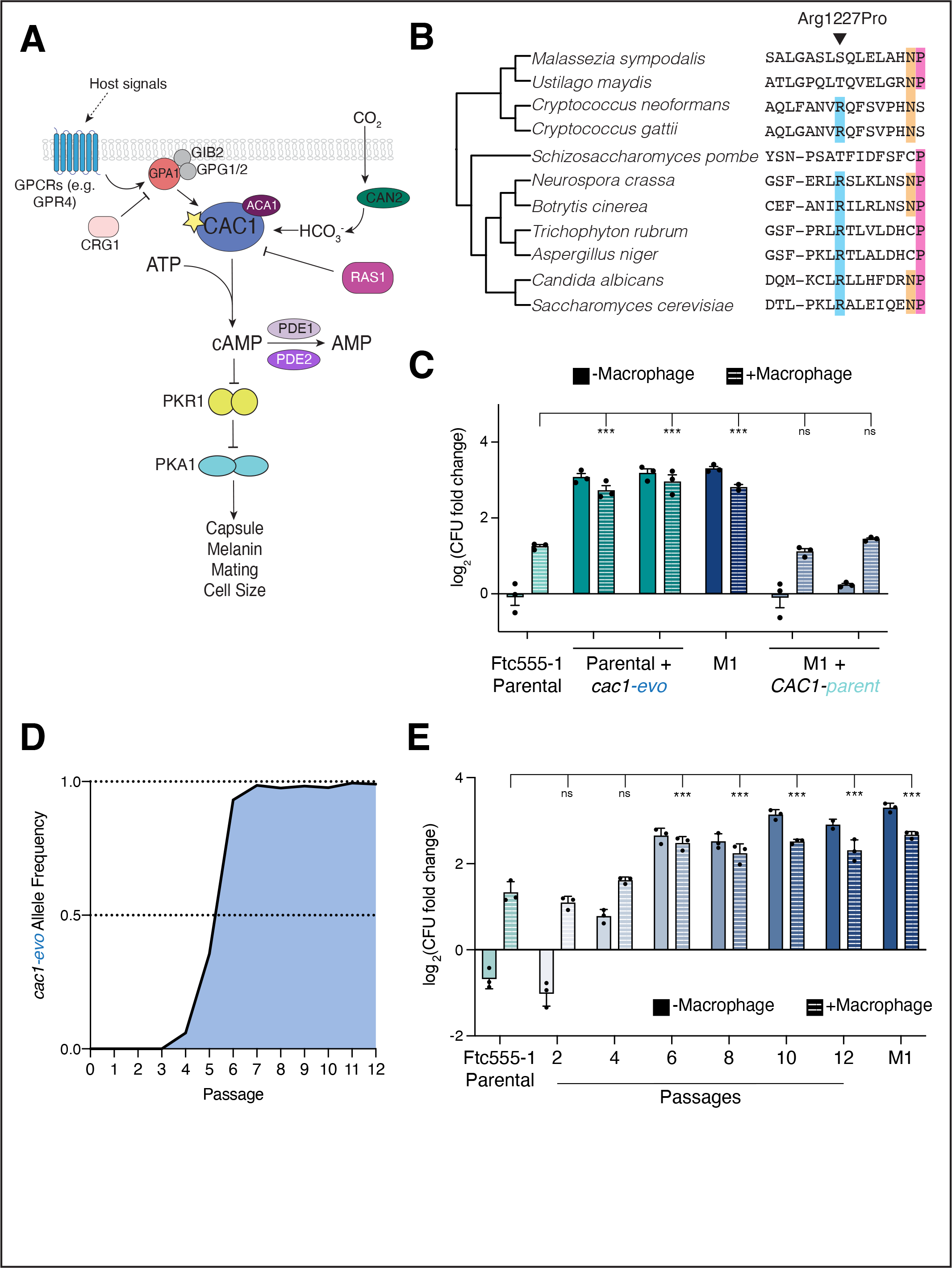
A single nucleotide mutation in the *CAC1* adenylate cyclase gene is necessary and sufficient for the enhanced growth of the evolved M1 strain. (A) Schematic of the cAMP signaling pathway in *C. neoformans*. *CAC1*, indicated with the yellow star, regulates the production of cAMP downstream of a number of host signals including nutrients and carbon dioxide. Production of cAMP leads to the activating of the PKA1 protein kinase which functions to regulate many key aspects of *C. neoformans* biology and virulence, including capsule and melanin production, mating, titan cell formation, and others. Activity of Cac1 and modulation of cAMP levels is achieved through the activity of a number of other genes, as indicated. (B) An arginine at amino acid position 1227 is conserved across fungi. Sequence alignments of the adenylate cyclase orthologs from a collection of fungal species; only the region surround the isolated mutation is shown. Position 1227, the site of the *cac1-evo* mutation, is indicated by the black arrow, and species with the conserved arginine at this site are shaded in blue. Other residues conserved among fungi are indicated in colors towards the 3’ end of this region. (C) Replication phenotypes of allele swap strains in macrophage co-culture experiments. Allele swap samples tested are two independently derived strains from CRISPR-mediated allele swap transformations (see Methods). (D) Allele frequency of the *cac1-evo* allele over the course of the evolution experiment. The number of reads containing the P1227R mutation in sequencing data from each passage population were counted and normalized to the read depth to calculate allele frequencies. (E) Growth of intermediate passage populations in co-incubation experiments with mouse macrophages. Cultures were started from frozen stocks of the passage populations (not individual colonies) and every other passage was tested. For (C) and (E), data shown is from one representative experiment with each dot representing a single replicate from that experiment. Solid bars show media-only growth, striped bars show growth in macrophages. Significance in these experiments was assessed compared to the parental strain by ordinary one-way ANOVA followed by Dunnett’s multiple comparisons test. *** p<0.0001, ns, not significant.

This mutation falls outside of the catalytic domain and other functionally annotated regions of Cac1, so its potential to alter Cac1 function and cAMP signaling is not obvious. To assess whether this position is conserved across *C. neoformans* or other fungi, we collected and aligned the protein sequences for adenylate cyclase orthologs from a sampling of other fungi, including both pathogenic and nonpathogenic species (Figure 4B). Although sequence conservation of these adenylate cyclase genes in fungi was generally quite low, we did observe conservation of an arginine at the corresponding amino acid position in most fungal species examined. Tellingly, in species where the arginine was not conserved, we see no evidence for substitutions as potentially disruptive as a proline at this site. Further, we looked for sequence variation at this position across the ~500 clinical and environmental isolates of *C. neoformans* that currently have sequence information available within the FungiDB repository. We saw 100% conservation of the arginine at amino acid position 1227 across all *CAC1* sequences in these isolates, though there was coding sequence variation in other regions of the gene. Together, the conservation of this site within *Cryptococcus* as well as across fungi is highly suggestive that this site is functionally important for the activity of Cac1.

The importance of Cac1 for *C. neoformans* biology combined with the lack of other mutations in the M1 strain were strongly suggestive that the Arg1227Pro mutation was likely causative for the adaptive phenotypes in this strain, which we sought to confirm experimentally. We swapped this single nucleotide position in both the parental Ftc555-1 and M1 strains using CRISPR-mediated gene editing (see Experimental Methods) and tested allele-swapped strains for growth in our macrophage co-culture experiments (Figure 4C). When the evolved *CAC1* allele— referred to from here on as *cac1-evo*—was introduced into the parental Ftc555-1 background, this was sufficient to cause a dramatic increase in the ability of this strain to replicate to high levels both in culture with macrophage cells as well as in tissue culture media alone. In the reciprocal swap, where the *cac1-evo* allele was replaced with the original parental version (*CAC1-parent*), we observed complete loss of the enhanced growth phenotypes. These data indicate that this single point mutation in the *CAC1* adenylate cyclase gene is both necessary and sufficient to confer the drastically altered phenotypes that we observe in our evolved M1 strain.

### Evolutionary dynamics of the CAC1 polymorphism reveal a strong competitive advantage associated with the evolved allele

We were curious about the emergence and dynamics of the *cac1-evo* allele over this short course of evolution. Initial sequencing revealed that this allele was fixed in the final population, but when did it arise and how quickly did fixation happen within the population? Taking advantage of the fossil record created during the evolution experiment, we extracted DNA from every frozen passage stock for whole genome sequencing and analyzed the *CAC1* locus in each population. By passage four a very low number of reads (ten total reads, allele frequency=~5%) contained the evolved allele, suggesting that the mutation arose *de novo* around this time (Figure 4D). The allele frequency of the *cac1-evo* allele rose quickly over subsequent passages: greater than 90% of reads contained the mutation at passage six and it had fully swept through the population by passage seven. This suggests that the *cac1-evo* allele confers a strong fitness advantage, in alignment with the competitive advantage we had observed in earlier experiments.

To examine the correlation between the *cac1-evo* allele and adaptation to macrophage host cells, we assessed the growth of intermediate passage populations in co-incubation experiments (Figure 4E). In agreement with the sequencing data, we see no significant difference in growth of the early passage populations. By passage four, growth in macrophages had trended upwards and by passage six, growth of the population in macrophages was indistinguishable from what we observed for the clonal M1 strain harvested from the end of the experiment.

We also assessed the competitive advantage of each passage population in head-to-head competition experiments with the NAT^R^ labeled parental strain (Supplemental Figure 4). We noted an increase in the competitive index for growth in macrophages beginning around passage six and reaching maximal levels at roughly passage 8, consistent with observations in tests of individual populations (Supplemental Figure 4A and Figure 4E). Media only competition experiments revealed a dramatic increase in competitive advantage beginning in passage five and which remained largely unchanged across subsequent passage populations (Supplemental Figure 4B). This largely agrees with our sequencing data and we hypothesize that the added complexity of the cultures in these competition experiments might explain the subtle discrepancies from the data we collected for the intermediate passages individually. Together, these results functionally link the emergence of macrophage adaptation in the M1 strain to mutation of *CAC1* and highlight the strength of the fitness advantage conferred by the *cac1-evo* allele, which rapidly swept through the population.

### *The* cac1-evo *allele causes changes in pathogenicity-associated traits*

cAMP signaling downstream of Cac1 is required for several critical biological functions important for the pathogenicity of *C. neoformans*^37^. For example, cAMP signaling is required for production of the characteristic polysaccharide capsule, which provides a protective outer coating and can shield cryptococcal cells from recognition by host phagocytes, including macrophages^32,34,35,38^. Cac1 activity has also been shown to be required for the production of melanin, another critical factor associated with enhanced survival of *C. neoformans* and other fungi *in vivo*^32,39^. To determine whether these downstream phenotypes are affected by the mutation we recovered in *CAC1*, we tested both melanin production and capsule formation in our parental and evolved strains. When plated onto melanin-inducing L-DOPA plates, there was no observable difference in the melanization of the parental or evolved strains of Ftc555-1 compared to each other or to the laboratory control strain KN99α (Figure 5B). Further, when we knocked out the full *CAC1* gene in both the Ftc555-1 parental and M1 background, we observed complete loss of melanin production, comparable to what has been observed for *cac1*Δ alleles in the laboratory strain background^32^. This suggests that while Cac1’s role in melanin production is conserved across these strains, the Arg1227Pro mutation likely does not confer a complete loss of function of Cac1 activity since melanization is preserved in strains carrying this allele.

**Figure 5.**
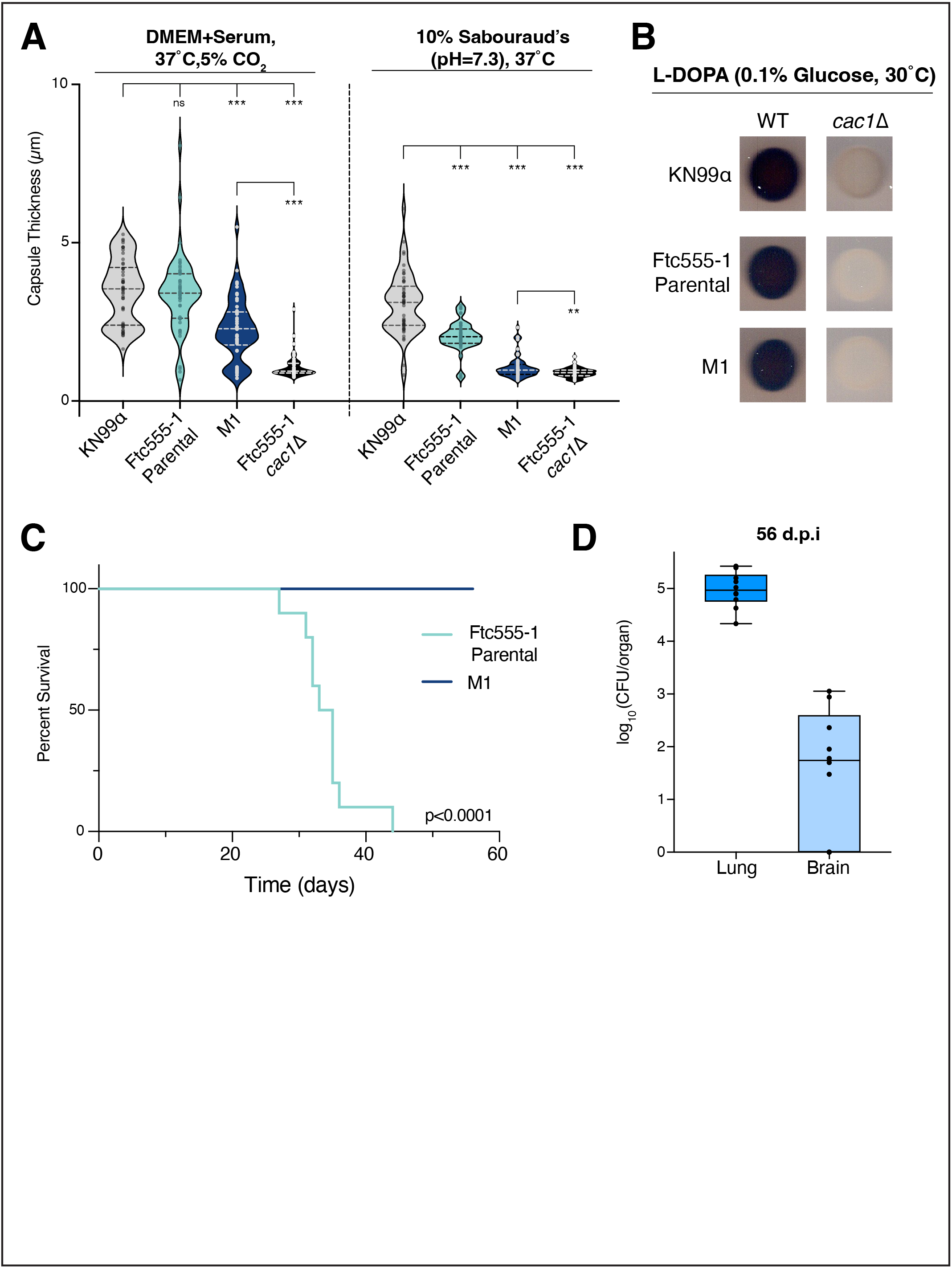
The *cac1-evo* allele affects pathogenicity-associated traits and decreases virulence in murine infection models. (A) Measurements of capsule size under two different capsule inducing conditions: tissue culture conditions (left side), or 10% Sab’s (right side). Violin plots show the distribution of measurements for 50 cells. Significance was assessed across all four tested strains by ordinary one-way ANOVA followed by Dunnett’s multiple comparisons test. Significance between M1 and the Ftc555-1 *cac1*Δ strain was assessed by an unpaired t-test with Welch’s correction. *** p<0.0001, ** p< 0.001, ns not significant. (B) Melanization of Ftc555-1 derived and control strains on L-DOPA solid media. The dark color of the strains indicates melanization (WT column). Deletion of the *CAC1* coding region in any of the strain backgrounds abolishes melanin production, indicated by the white color of these spots on L-DOPA (*cac1*Δ column). (C) Survival of C57BL/6NJ mice following intranasal inoculation with Ftc555-1 (teal) or M1 (navy blue). Ten animals were inoculated per strain (five male, five female) and were weighed daily until they reached predetermined endpoint criteria. Significance was determined by a log-rank test. (D) Fungal burden in the lung and brains of mice infected with the M1 strain from (C) on day 56 post infection. Each data point indicates the CFU count from an individual animal.

Next, we looked at capsule size in our parental and evolved strains. Capsule is induced by specific culturing conditions that mimic host-like environments, which include: elevated pH, nutrient limitation, high temperature and increased CO_2_ concentration^40–42^. We tested the ability of our strains to induce capsule using two different induction protocols (see Experimental Methods); mutants in the cAMP signaling pathway have previously been suggested to be sensitive to the specific capsule induction conditions, motivating our decision to test multiple protocols^41^.

We observed both strain-specific and condition-specific differences in capsule induction in the Ftc555-1 parental and M1 evolved strains. Across both conditions, cells from the parental strain produced significantly larger capsules than M1 cells (Figure 5A). Under tissue culture conditions (DMEM+Serum+CO_2_), the parental Ftc555-1 cells had capsule sizes comparable to the laboratory control strain, KN99α, but capsule size in this strain was significantly reduced in the alternate capsule inducing protocol (10% Sab’s). M1 cells did produce capsule under tissue culture conditions, albeit at a reduced size compared to other strains. However, capsule production was markedly reduced in M1 cells incubated in the Sab’s inducing media. M1 capsule sizes under these conditions dropped nearly down to the levels of an acapsular *cac1*Δ mutant (Figure 5A). In contrast to the Ftc555-1-derived strains, we observed no significant differences in capsule size across different conditions for the control strain, KN99α. Similarly, a *cac1*Δ mutant in the Ftc555-1 background produced very little capsule in either condition. This confirms that Cac1 activity is required for capsule production in the Ftc555-1 genetic context as it is in laboratory reference strains^32^. Collectively, these data suggest that *cac1-evo* encodes a partial loss of function allele as evidenced by reduced capsule production in the M1 strain. However, the dependence of capsule phenotypes on the conditions used hint at a more complex mechanism underlying capsule regulation, perhaps reflecting different activity requirements for the Cac1 pathway under distinct nutritional and environmental conditions.

### *The* cac1-evo *allele causes decreased mortality in murine infection*

Mutation of genes in the cAMP signaling pathway, including *CAC1*, have been reported to be associated with substantial changes in *in vivo* phenotypes^32,34,35^. We used a murine model of disseminated cryptococcosis by inoculating C57BL/6N mice intranasally with either Ftc555-1 or the derived M1 strain to assess the pathogenicity of these strains. Our *in vitro* results comparing parental and evolved strains of Ftc555-1 predicted two possible and opposite outcomes for the pathogenicity of these strains *in vivo.* The competitive advantages in intracellular macrophage replication we observe in our evolved strain suggest a model where the M1 strain might similarly replicate and disseminate to higher levels in the context of an animal infection. In contrast, the capsule size differences observed for the M1 strain, predict a potential tradeoff involving decreased fitness for this strain *in vivo* due to enhanced recognition and control by the immune system. Consistent with the second possibility, we observed a striking difference in survival between the two strains in our animal model of infection (Figure 5C). Animals infected with the parental strain, Ftc555-1, had a median time to endpoint of 34 days, while the M1 strain failed to cause lethal infection in any animals during the experiment.

To address whether the lack of disease in animals infected with the M1 strain was due to clearance of *C. neoformans* after inoculation, we sacrificed animals 56 days after infection and assessed fungal burden by plating homogenized lung and brain tissue for CFU counts (Figure 5D). Intriguingly, we recovered CFUs from both the lungs and the brains of M1-infected mice; CFU levels in the lungs were, on average, higher than the starting inoculum (>10^5^ CFU/lung). CFUs in the brain were low, though we observed dissemination to the brain to varying extents in seven out of ten of the infected animals. These results indicate that mice were persistently infected with *C. neoformans* and that the M1 strain maintains its potential for dissemination. Although seemingly paradoxical given fitness improvements observed in our *in vitro* macrophage experiments, these results show the remarkable effect a single nucleotide change in the genome of a pathogen can have on the disease-causing potential of the organism. This surprising tradeoff in adaptation to macrophages leading to decreased pathogenicity in mouse infections raises new questions about the impact different environments play in shaping the evolution of fungal pathogens.

## Discussion

The role of evolutionarily diverse hosts in influencing the acquisition and evolution of pathogenicity among infectious microbes is a fundamental and fascinating question. Here, we harnessed naturally occurring genetic variation in the fungal pathogen *C. neoformans* to investigate the evolutionary mechanisms underlying host adaptation and its relationship to pathogenicity in this species.

An emerging body of work examining strain-level variation in both *in vitro* and *in vivo* assays has demonstrated that even closely related *C. neoformans* strains can have divergent phenotypes in these experiments^4,5,8–10,43,44^. Our study provides a major advance to this field by directly comparing the growth of the same panel of clinical and environmental isolates in host cells from two different species. We identified strain-specific differences in survival during interactions with the amoeba species, *A. castellanii*, or mouse macrophage-like cells across this panel of strains (Figure 1B-E). In particular, we noted that there was correlation between strains that were able to withstand killing by amoebae and those that replicated the most in macrophages. This suggests that there may be similarities in the intracellular environments of these diverse host species that favor the growth of certain *C. neoformans* isolates over others. Future work comparing genome sequences of these strains to reveal the genetic basis of these phenotypes will help to shed light on commonalities between host environments that could lead to the emergence of pathogenic (or more pathogenic) strains. Our work also highlights the importance of comparative studies, looking at survival of *C. neoformans* isolates in a range of different host species in revealing additional adaptive strategies that confer widespread evolutionary success for these microbess both in patients and in the environment.

Our courses of experimental evolution revealed that adaptation to host conditions can happen over extremely short timescales in *C. neoformans* (Figure 2). Rapid adaptation of pathogens to new hosts or new environments is well documented for viral and bacterial pathogens^45–48^. But most previous studies of host adaptation in pathogenic fungi, have focused on evolution occurring over much longer timescales^25–27,49^. We estimate that the macrophage-passaged lines generated here were evolved for roughly 75-100 total generations including outgrowth, while amoeba-passaged strains likely experienced significantly fewer generations due to their potent killing by amoeba in early passages. But while rapid adaptation was observed in some of the serially passaged strains, not all strains had noticeable changes in phenotype, highlighting the power of these serial passaging approaches to reveal a range of evolutionary trajectories across different replicates.

These experiments also revealed adaptive phenotypes that are likely host-specific, conferring an advantage in either amoebae hosts or in mammalian macrophages, but not in both. Although some virulence-associated traits of *C. neoformans* have been shown to be important for growth in many different cell types, the extent to which intracellular survival mechanisms are shared across hosts is currently unknown^19,20,28,50^. The isolates we recovered here, along with future iterations of this experimental evolution, will help shed light on conservation and divergence in host selective pressures on *C. neoformans* adaptive evolution.

Characterization of one of our host-adapted strains—macrophage-passaged line M1— revealed that the acquisition of a single point mutation in the adenylate cyclase gene *CAC1* conferred a strong competitive advantage for growth in macrophages. The effect of this mutation on Cac1 function and the mechanism by which this contributes to the growth phenotypes we observe in M1 is an exciting open question. Our data on downstream phenotypes controlled by the cAMP signaling pathway revealed that this mutation does not abolish Cac1 function entirely, as strains carrying the evolved *cac1-evo* allele remain competent to produce melanin and limited amounts of capsule. Our results on capsule size across different conditions suggest that Cac1 activity may be differentially regulated by different environmental and/or nutrient cues and that the evolved allele may be particularly responsive to such fluctuations in growth conditions. We hypothesize that the Arg1227Pro mutation might affect interactions between Cac1 and regulatory proteins that help to control Cac1 activity under these different conditions, such as Gpa1, Aca1 or Ras1. Future genetic and biochemical analyses of this allele will explore the complex impact of this point mutation in *CAC1*.

It is also important to note that previous studies of cAMP signaling generally and Cac1 specifically, were carried out almost exclusively in the context of the laboratory strains H99 and KN99. One notable exception has been the study of PKR1 variation downstream of *CAC1* and its role in titan cell formation in clinical isolates of *C. neoformans*^44^. Our analyses of *CAC1* sequences from diverse *C. neoformans* isolates revealed that there was considerable sequence variation in this gene across this species. In the Ftc555-1 parental strain alone, there are 17 coding changes in *CAC1* compared to the most commonly studied H99 and KN99 lineage strains. This suggests that there may be functionally significant differences in the activity of this critical signaling pathway due to sequence changes in *CAC1* or through epistasis with other genes, like *PKR1*, with altered sequences in these variant strains. Curiously, *PKR1* mutations have also been identified in serially sampled isolates from human patients, as well as in amoebae-adapted strains, though the functional significance of these mutations has not been fully investigated ^24,50^. Regardless, the convergence of multiple studies, including ours, on the cAMP signaling pathway suggest it may be a hotspot of genetic variation within this species, with functional implications for a range of pathogenicity-associated phenotypes. This further underscores the power of using naturally occurring genetic variation found in clinical and environmental isolates of *C. neoformans* to gain new insight into key virulence pathways that have previously only been studied in a single genetic context.

Finally, our results raise intriguing questions about the relative tradeoffs between adaptation to specific host cell environments and maintenance of pathogenicity, or the ability to cause disease. Our *in vivo* experiments revealed a striking difference in the pathogenicity of the serially passaged M1 strain when compared to the parental Ftc555-1 strain. Given its markedly enhanced ability to replicate in mouse macrophages, we were surprised that M1 failed to cause lethal disease in a mouse model. Many *in vitro* phenotypes of improved pathogen replication are well correlated with more severe disease outcomes in hosts. Whether the lack of disease we observe in M1-infected animals is due to an altered host immune response or an inability of this strain to replicate well in the context of the more complex environment of the mouse lung is an open question. Future work looking at host inflammatory responses, differences in lung physiology and disease progression during infection with this strain will help distinguish between these possibilities. Regardless of exact mechanism, the complex impact of a single nucleotide change in the genome of this pathogenic fungus highlights the long reach of simple adaptive changes to fundamentally change the course of evolution.

## Methods

### Strains and growth conditions

*C. neoformans* strains used in this study are listed in Supplementary Table 1. Strains were stored at −80°C in yeast extract peptone dextrose (YPD) media supplemented with 20% glycerol. Strains were inoculated by streaking onto YPD plates and incubated for 2-3 days at 30°C before use in experiments. Strains were stored at 4°C for <10 days and reinoculated as needed. Unless otherwise noted, all overnight cultures were grown for 16-18 hours in YPD at 30°C with shaking at 225 rpm.

### *Macrophage and* A. castellanii *cell culture*

Both J774A.1 cells and *Acanthamoeba castellanii* were purchased from ATCC. The J774A. 1 cell line was cultured at 37°C with 5% CO_2_ in DMEM with high glucose and L-glutamine (VWR 16777-129) supplemented with 10% heat inactivated FBS, non-essential amino acids and penicillin-streptomycin. The *A. castellanii* strain ATCC 30234 was cultured axenically in Peptone-Yeast-Glucose (PYG) Media (2% proteose peptone, 0.1% yeast extract, 0.1 M glucose, 4 mM MgSO4, 0.4 M CaCl_2_, 0.1% sodium citrate dihydrate, 0.05 mM Fe(NH_4_)_2_(SO_4_)_2_• 6H_2_0, 2.5 mM NaH_2_PO_3_, 2.5 mM K_2_HPO_3_, pH 6.5) in T175 tissue culture flasks at 28°C. Both cell types were used for assays between passages 5 and 20 from thaw.

### Co-culture experiments with macrophages and amoeba

Macrophages were seeded into two replicate 96-well plates 18-24 hours before each assay at a density of 10^4^ cells/well in 100 μL normal macrophage growth media and incubated at 37°C and 5% CO_2_. Overnight cultures of each *C. neoformans* strains were subcultured down to an OD of 0.2 and allowed to reach mid-log phase (4-5 hours, OD=0.6-1.0) before use in experiments. Each strain to be tested was then washed twice with sterile 1x PBS and adjusted to a concentration of ~2×10^5^ cells/mL in macrophage growth media containing 1 μg/mL of the anti-GXM antibody 18B7 (Sigma Cat# MABF2069). Cells were opsonized in this media for 1h at 37°C. While *C. neoformans* cells were opsonizing, macrophages were activated for 1 hour by replacing the media with 100 μL of growth media with 10 nM phorbol myristate acetate (PMA). Media was removed from activated macrophages and 100 μL of opsonized *C. neoformans* suspensions were added to each well (2×10^4^ yeast cells/well for an MOI ~1). 100 μL of opsonized cell suspensions were also added to empty wells in the plate as controls for media growth. *C. neoformans* cells were allowed to be phagocytosed for 1 hour with incubation at 37°C with 5% CO_2_. After 1 hour, supernatant and extracellular yeasts were removed and macrophages were gently washed 1-2 times with warm PBS before replacing with fresh macrophage growth media. One replicate plate was immediately lysed with 200 μL of 0.1% sodium deoxycholate to determine starting CFU and percent phagocytosis counts. Wells were washed twice with sterile water to collect all *C. neoformans* cells and combined with lysates. The remaining plate was incubated for an additional 24 hours before lysis. Lysed cultures were serially diluted and plated onto YPD. Plates were incubated at 30°C for 2-3 days before CFUs were enumerated.

Fold change in CFU was calculated by comparing the CFU counts after 1 hour of phagocytosis to the 24 hour CFU counts. Media only fold changes were calculated by comparing CFU counts from the media only wells at 1 hour vs. the media only wells at 24 hours. Percent phagocytosis was determined by comparing the 1 hour CFU counts from wells with macrophages to the control wells with media only.

Amoeba co-culture experiments were carried out similarly to the macrophage experiments with a few modifications. Amoeba were seeded at a density of 5×10^4^ cells/well into replicate 96-well plates and *C. neoformans* cell suspensions were prepared at 1×10^6^ cells/mL in Ac Buffer (4 mM MgSO4, 0.4 M CaCl_2_, 0.1% sodium citrate dihydrate, 0.05 mM Fe(NH_4_)_2_(SO_4_)_2_• 6H_2_0, 2.5 mM NaH_2_PO_3_, 2.5 mM K_2_HPO_3_, pH 6.5) to maintain an MOI of ~1. Amoeba were starved of nutrients in Ac Buffer for 1 hour at 28°C before 100 μL of yeast cell suspensions were added, and the cultures remained in Ac Buffer throughout the duration of the experiment. All incubations were performed at 28°C Timings, washes, and CFU enumeration was carried out as described above. For all co-incubation experiments, each condition and strain was tested in triplicate and experiments were repeated 2-3 times on different days unless otherwise noted.

### *Serial passaging of* C. neoformans *through host cells*

8.5-9 × 10^5^ macrophage or *A. castellanii* cells were seeded into T25 flasks 16-18 hours before each passage and incubated under their respective normal growth conditions. For the first passage, an overnight culture of Ftc555-1 was grown with shaking for 16 hours with shaking at 30°C. This culture was centrifuged at 3000 × g for 5 minutes and washed twice with 1x PBS before the OD was measured. Cells were normalized to an OD of 1 (~1.5 × 10^7^ cells/mL) in 3.5 mL of either macrophage growth media containing 1 μg/mL 18B7 anti-capsule antibody (for macrophage passaging) or in Ac Buffer (for amoeba passaging). Cell suspensions were incubated at 37°C (macrophage passaged) or 28°C (amoeba passaged) for 1 hour. The remaining starting inoculum culture was mixed 1:1 with 40% glycerol and aliquots were frozen at −80°C.

While *C. neoformans* cells were incubating, the media from macrophage containing flasks was removed and replaced with macrophage growth media containing 10 nM PMA. PYG media was removed from amoeba flasks and replaced with Ac buffer and cells were incubated at their standard temperatures for 1 hour. Following this 1 hour incubation, 1 mL of the appropriate *C. neoformans* cell suspension was added to each flask and gently rocked to ensure the entire host cell layer was covered. This small volume ensured that yeast cells settled onto the host cell monolayer and facilitated phagocytosis. Phagocytosis was allowed to proceed for 2 hours before extracellular yeasts were removed. Cells were washed 1-2 times with pre-warmed PBS and media was replaced with 2 mL of growth media. Phagocytosed yeasts and host cells were incubated for 24 hours.

After 24 hours of incubation, the media containing extracellular yeasts was discarded from lines 1 and 2 of both amoeba- and macrophage-containing flasks. One mL of 0.1% sodium deoxycholate was added to all of the flasks to lyse host cells and lysates were collected. Flasks were washed twice with 1x PBS. Lysates and washes were combined and centrifuged at 3000 × g for 5 min and the supernatant was discarded. *C. neoformans* cell pellets were resuspended in YPD media, vortexed to mix, and incubated at 30°C with shaking (225 rpm) for 24 hours.

Following outgrowth, ODs were measured and each individual culture was washed and adjusted to an OD of 0.425 in 1.5 mL of the appropriate growth media. The remaining culture was frozen as above to create a fossil record for the experiment. Passaging proceeded as above from this point. All macrophage passages were performed at 37°C with 5% CO_2_. Amoeba passages were largely carried out at 28°C, but passages 5 and 10 were performed at 37°C with 5% CO_2_ to avoid losing thermotolerance in these strains. All outgrowth steps were carried out at 30°C. Following the outgrowth of the final passage, multiple aliquots were frozen. Samples from each population were also serially diluted and plated onto YPD plates for CFU determination. Individual colonies from these plates were picked into 5 mL of YPD, grown overnight and frozen for use in phenotyping experiments.

### Strain construction and molecular biology

Primers used in this work are listed in Supplemental File 2. All genetic manipulations were done using CRISPR-mediated gene editing and the TRACE system following previously published protocols^51,52^. Briefly, gRNAs were generated using guide-specific primers and amplification from the pBHM2329 plasmid (gift from H. Madhani). *CAS9* was amplified for transformation from the pBHM2403 plasmid. All DNA used for CRISPR transformations was generated by PCR and purified using the Zymo DNA Clean and Concentrator 25 kit. If necessary, ethanol precipitations were performed to further concentrate DNA. Cells were grown as described for transformations and DNA was introduced by electroporation with an Eppendorf Eporator at 2 kV. DNA concentrations used were as follows, unless otherwise noted: 250 ng Cas9, 100 ng gRNA, 2.5 μg Repair Template.

For insertion of a the nourseothricin cassette at the <u>S</u>afe <u>H</u>aven 2 locus (SH2), we designed a VNB lineage-specific guide targeting this locus^53^. A repair template plasmid carrying the ~1 kb of sequence immediately 5’ and 3’ to this gRNA sequence from the H99 genome was constructed using HiFi DNA Master Mix (NEB Cat#E2621) with a standard NAT^R^ cassette inserted between the two flanks. Repair templates were then PCR amplified from this plasmid for CRISPR transformations.

For allele swap strains, the M1-derived mutation in *CAC1* conveniently generates a new PAM site in this strain, so we used strain-specific gRNAs that overlap or terminate at this mutation, wherein the PAM and guide sequences would then be disrupted by introduction of the alternate allele. Repair constructs containing the site of the mutation and ~900 bp upstream and downstream of this position were amplified directly from genomic DNA from either Ftc555-1 or M1. We used a co-CRISPR approach, simultaneously targeting *CAC1* and the SH2 locus in order to be able to select for successful transformation. For allele swap transformations, we simultaneously transformed the following amounts of DNA: 250 ng M1- or Ftc555-1-specific gRNA, 250 ng VNB SH2 gRNA, 1 μg Cas9, 2 μg *CAC1* gDNA Repair Template, 2 μg SH2-NAT Repair Template.

For generating *cac1*Δ mutants in the Ftc555-1 and M1 backgrounds, gRNAs targeting the 5’ and 3’ ends of the gene were generated and both were used in transformations to ensure deletion of the entire open-reading frame. A repair template plasmid was created by amplifying the ~1kb regions upstream and downstream of the ORF of *CAC1* from Ftc555-1 gDNA. HiFi DNA master mix was used to insert a hygromycin resistance cassette (HYG^R^) between these flanking gDNA sequences and ligate it into the pUC19 background. Repair template was amplified from the resulting plasmid for use in CRISPR transformations.

### Transformant selection and genotyping

Following electroporation of CRISPR reagents, strains were plated onto selective media (either YPAD+NAT or YPAD+HYG) and allowed to grow for 2-3 days at 30°C. Individual colonies were then patched onto non-selective YPD media for 2-3 passages before being patched back onto selective media in order to identify stable genomic integrants.

Colonies were genotyped by colony PCR: small amounts of cells were picked into PCR strip tubes and microwaved for 2.5 minutes before a Phusion-Flash (Thermo Cat# F548S) PCR mix containing the appropriate primers was added directly to these cells. For SH2 and *cac1*Δ genotyping, a primer sitting inside of the inserted drug cassette on both 5’ and 3’ ends (ZAH_G40 and G01) were paired with primers in the flanking genomic regions outside of the repair template constructs (for SH2: MS_B01 and B02, for *cac1*Δ: ZAH_I06 and I07). Allele swap colonies were genotyped using primers 5’ and 3’ to the repair construct (ZAH_H20 and H21) paired with destabilized primers designed with the WebSNAPER tool to bind specifically to either the parental or evolved *CAC1* allele (ZAH_H22 for the parental allele and ZAH_H30 for the evolved allele)^54^. Allele swap colonies were first genotyped for the correct *CAC1* allele, and were then confirmed to also carry the NAT^R^ cassette at the SH2 locus.

Genomic sequences were amplified from gDNA extracted from candidate colonies and integration and allele identity was confirmed by Sanger sequencing. PCRs were also performed to ensure that no strains had integrated the gRNA or *CAS9* constructs.

### Competition experiments

For competition experiments, cell suspensions were made identically for test strains and the parental Ftc555-1 carrying a NAT-resistance cassette at the SH2 locus. These cell suspensions were mixed 1:1 before adding 100 μL to wells for co-culture experiments, which were carried out as described above. To determine the CFUs for test strain vs. the parental strain, serial dilutions were plated onto both YPD plates as well as YPAD+NAT plates. Competitive index here is calculated as:

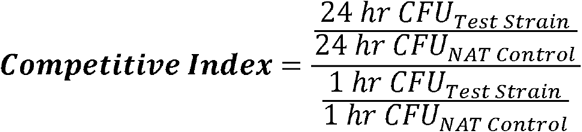

### Intracellular and extracellular population determinations

To determine intracellular and extracellular CFU counts, co-incubation experiments were set up as described above. After 24 hours of incubation with macrophages, the media supernatant containing extracellular yeasts was removed to an empty well of the 96-well plate and replaced with fresh macrophage media. If there were still substantial unassociated yeasts observed, this step was repeated for a second time. Otherwise, macrophages were lysed as above to collect intracellular yeasts. Both intracellular and extracellular wells were washed, serially diluted, and plated for CFU enumeration as described above.

### Fluconazole MIC determinations and competition experiments

Overnight cultures of *C. neoformans* strains were adjusted to a density of ~5 × 10^6^ cells/mL and were spread onto either YPD-agar or RPMI agar medium with L-glutamine and without sodium bicarbonate. Fluconazole Etest strips (bioMérieux) were placed on the agar surface of the RPMI plates once cultures had dried. All plates were placed at 37°C and imaged after 48 and 72 hours. Only strains with robust growth on control YPD plates were imaged and used to determine MIC values.

For macrophage experiments and competitions with fluconazole supplementation, co-incubation setup was performed as above through the one hour phagocytosis step. When extracellular yeasts and media were removed, media was replaced with standard macrophage growth media containing either 2 μg/mL Fluconazole (in DMSO) or media with an equal volume of DMSO. The remainder of the experiment was performed as described above.

### Genomic DNA extraction and sequencing

Single colonies or 250 μL of pooled glycerol stocks for sequencing were inoculated into 50 mL of YPD and grown overnight with shaking at 30°C. Cells were collected by centrifugation and lyophilized overnight. High-molecular-weight genomic DNA was isolated following the CTAB extraction protocol as previously described. Following CTAB extraction, samples were further purified using the Takara ChromaSpin 1000 column. Concentrations were measured by Qubit broad range assay. For Nanopore sequencing of Ftc555-1, one microgram of HMW gDNA was prepared for sequencing using the SQK-LSK110 Kit and run on an R9 (FLO-MIN106) flow cell. For Illumina sequencing, samples were sheared with a Covaris S2 Focused-ultrasonicator and libraries were prepared using the New England Biolabs NEBNext Ultra II DNA Library Prep kit (cat#E7634L) with an average insert size of 450 bp. 12-14 strains per run were barcoded and pooled, and 100 million 150×150 bp paired-end reads were collected on a NovaSeq S4 Flow Cell with the v1.5 Reagent Kit. Illumina library preparation and sequencing was performed by the University of Utah’s High Throughput Genomics core facility at the Huntsman Cancer Institute.

### Sequencing analysis

Fast5 files from Nanopore sequencing of the parental strain were re-basecalled using guppy-gpu (v.6.0.1). The resulting fastq files were directly used for assembly using Canu (v 2.1.1) with an estimated genome size of 20 Megabases and default parameters^55^. This assembly had 14 chromosome-scale contigs and a number of shorter contigs which mapped either to telomeric regions or mitochondrial DNA. These were manually trimmed from the assembly. The assembly was polished once with Nanopore reads using Medaka (1.5.0; https://github.com/nanoporetech/medaka) and then polished five times with Illumina reads from the parental strain using pilon (v 1.24) until no additional changes were being made to the assembly^56^. A circular mitochondrial genome was assembled from Illumina reads using GetOrganelle (v1.7.5.3) with the fungus_mt setting and confirmed by comparison to the trimmed contigs assembled by Canu^57^. This mitochondrial genome was manually added to the assembly. Chromosomes were named based on comparison to other reference assemblies and annotations were transferred using LiftOff (v1.6.3)^58^.

For Illumina sequencing data of evolved colonies and passages, adapters were trimmed using Trimmomatic (v 0.39) and reads were aligned to our *de novo* assembly using bwa-mem2 (v 2.2.1)^59,60^. Duplicates were marked and read groups added with Picard and the resulting files were sorted and indexed with SAMtools^61^. Variants were called using sorted BAM files with the Genome Analysis Tool Kit (HaplotypeCaller and GenotypeGVCF, v3.8) with the ploidy set to 1 ^62^. Variants were filtered out based on the following criteria using GATK VariantFiltration: QD<2.0, FS > 60, MQ <40, GQ <50, DP <10. To remove called SNPs in homopolymeric runs likely caused by sequencing errors, we used the BCFtools isec function to identify unique SNPs in each sample^61^. The predicted impacts of called variants were assessed using SnpEff using our *de novo* assembly as reference^63^.

To analyze allele frequencies of the *cac1-evo* allele, sorted BAM files from each passage were visualized in IGV and the number of reads containing the Arg1227Pro (Chromosome 8: 1057343 G>C) were counted and normalized to the total sequencing coverage at that position^64^. All sequencing analysis was performed using resources through the University of Utah Center for High Performance Computing.

### Melanization and capsule assays

Melanization of different strains was analyzed following previously established protocols^65^. Briefly, overnight cultures of *C. neoformans* strains were pelleted by centrifugation (3000xg, 5 minutes), washed twice in sterile water and adjusted to an OD of 0.25. Five microliters of each strain was spotted onto L-DOPA plates (7.6 mM L-asparagine monohydrate, 5.6 mM glucose, 10 mM MgSO_4_, 0.5 mM 3,4-dihydroxy-L-phenylalanine, 0.3 mM thiamine-hydrochloride, and 20 nM biotin) and onto YPD plates and incubated at 30°C. Plates were checked every day until the WT control (KN99α) turned dark brown, usually day 3. The plates were then imaged using an Epson Perfection V39 scanner.

Capsule was induced using two different protocols^40^. For both protocols, *C. neoformans* strains of interest were grown overnight (16-18 hours) in YPD media and OD_600_ was measured for each strain. In the first protocol, strains were subcultured to an OD of 0.3 in DMEM media containing 10% FBS, non-essential amino acids, and penicillin/streptomycin and incubated in a 6-well plate at 37°C with 5% CO_2_. In parallel, the same subcultures were subcultured to an OD of 0.3 in 10% Sabouraud’s Dextrose Media with 50 mM HEPES adjusted to pH 7.4. Strains in 10% Sab’s were incubated in 6 well plates at 37°C without CO_2_. As a control, the same strains were diluted in YPD media to the same OD and replicate plates were incubated at 37°C with or without CO_2_. Cell and capsule size measurements were made after 48 hours of incubation. Cells were centrifuged at 3000 x g, resuspended in 50 μL of PBS and mixed with india ink to visualize capsule. 5-10 μL was pipetted onto a microscope slide and pictures were taken on an EVOS FL Cell Imaging System (Life Technologies/Thermo Fisher) at 40x magnification. For each strain, measurements were made for 50 cells under each condition using FIJI^66^. Capsule thickness here was calculated as:

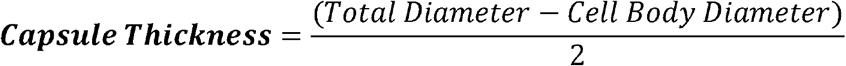

### Animal experiments

~8-week-old C57BL/6NJ mice were ordered from Jackson labs and randomly assigned to experimental groups, with 5 male and 5 female mice tested per condition. Mice were anesthetized with ketamine/dexmedetomidine hydrochloride (Dexdomitor) delivered intraperitoneally. They were then suspended by their front incisors on a horizontal strand of thread. Mice were inoculated intranasally with 2.5×10^4^ *Cryptococcus* cells in 50 μl of 1XPBS using a micropipette. The inoculum was placed dropwise onto a nasal flare before being inhaled by mice. Ten minutes later, mice were intraperitoneally administered the reversal agent atipamezole (Antisedan). For survival analyses, mice were weighed daily and euthanized by CO_2_ asphyxiation and cervical dislocation when they lost 15% of their initial mass.

For fungal burden determination, the M1-infected mice were euthanized by CO_2_ asphyxiation and cervical dislocation at 56 dpi. Organs were harvested, placed on ice, and homogenized in 5 ml of 1x PBS. Serial dilutions of homogenates were plated on Sabouraud’s dextrose agar with 10 mg/mL gentamicin and 100 mg/mL carbenicillin. Plates were incubated at 30°C for 2-3 days before CFUs were counted to determine fungal burden per organ. Animal procedures were approved by the University of Utah Institutional Animal Care and Use Committee.

## Acknowledgements

We thank John Perfect for providing the clinical and environmental isolates used in this study and Hiten Madhani for providing plasmids for CRISPR transformations. We thank members of the Elde and Brown labs for helpful discussions in the development of this project, Kyla Ost for a lot of guidance and feedback and Vikas Yadav for help with DNA extraction and Nanopore sequencing protocols. This work was funded by a Helen Hay Whitney Foundation postdoctoral fellowship to Z.A.H., NIH T32 funding (T32AI055434) to K.Y.C and J.M.B, an NIH R01 (R01AI130248) awarded to J.C.S.B, a Burroughs Wellcome Fund Investigator in the Pathogenesis of Infectious Disease award to N.C.E., and an NIH R35 (R35GM134936) awarded to N.C.E.

## Author Contributions

Conceptualization: Z.A.H.

Methodology: Z.A.H., K.Y.C., J.M.B.

Software: Z.A.H.

Validation: Z.A.H.

Formal Analysis: Z.A.H.

Investigation: Z.A.H., K.Y.C., J.M.B., M.W.S.

Resources: J.C.S.B., N.C.E.

Data Curation: Z.A.H., K.Y.C., J.M.B.

Writing-Original Draft: Z.A.H.

Writing-Review and Editing: Z.A.H., N.C.E.

Visualization: Z.A.H.

Supervision: J.C.S.B., N.C.E.

Funding acquisition: J.C.S.B., N.C.E.

## Supplemental Figures

**Supplemental Figure 1.**
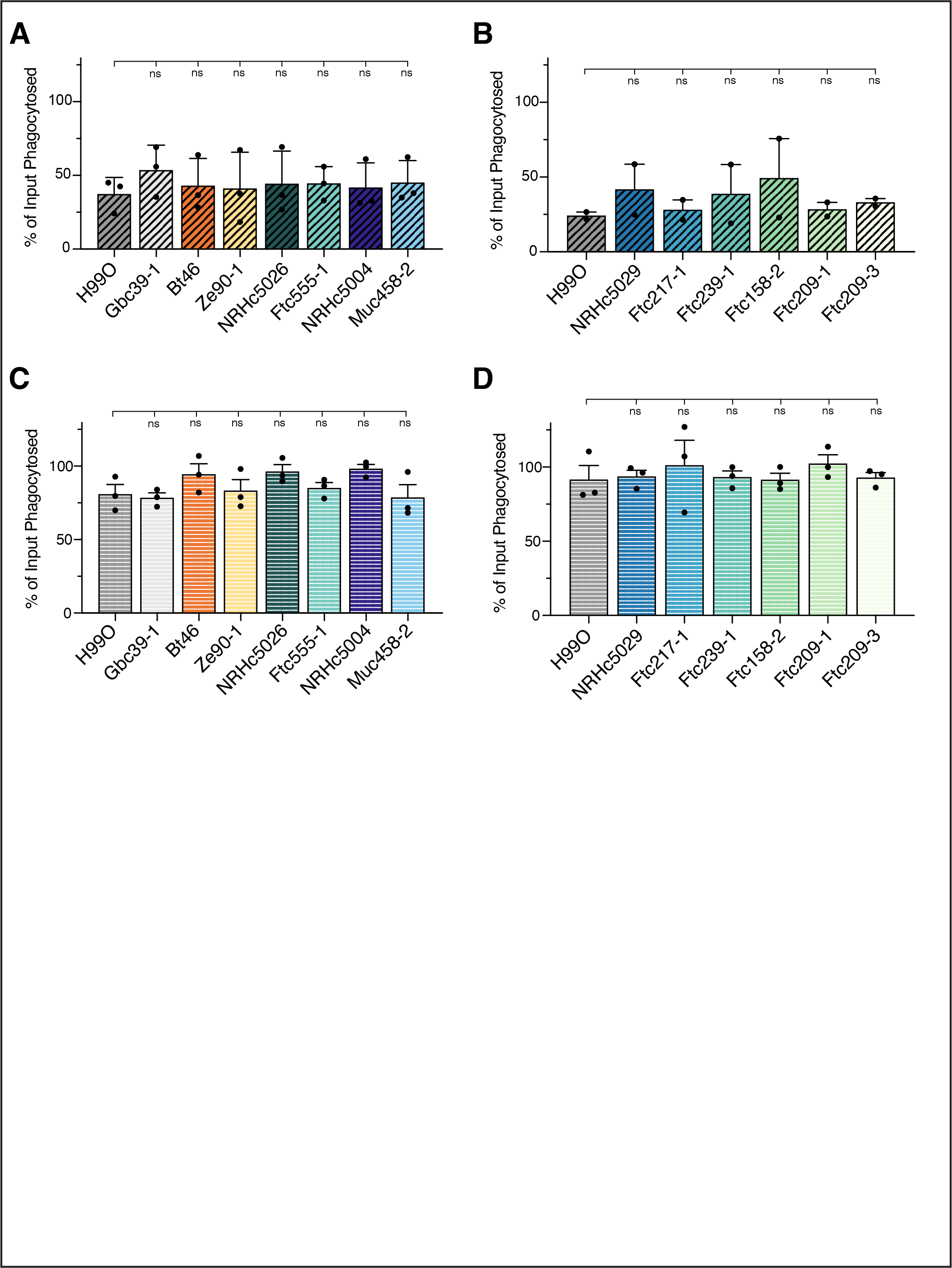
Differences in phagocytosis do not explain strain variation in host cell replication. The percent of cells from the input culture that were phagocytosed over one hour of incubation with *A. castellanii* cells (A and B) or mouse macrophages (C and D) was calculated as the number of CFUs recovered from wells containing host cells after 1 hour of co-incubation divided by the number of CFUs from control wells without host cells present. Plotted data indicate the average values ± SEM from 2-3 independent experiments on different days. Each dot indicates the average of three replicate measurements from a single experiment. Significance was assessed by comparison to the laboratory strain H99O using ordinary one-way ANOVA followed by Dunnett’s multiple comparisons test. ns, not significant.

**Supplemental Figure 2.**
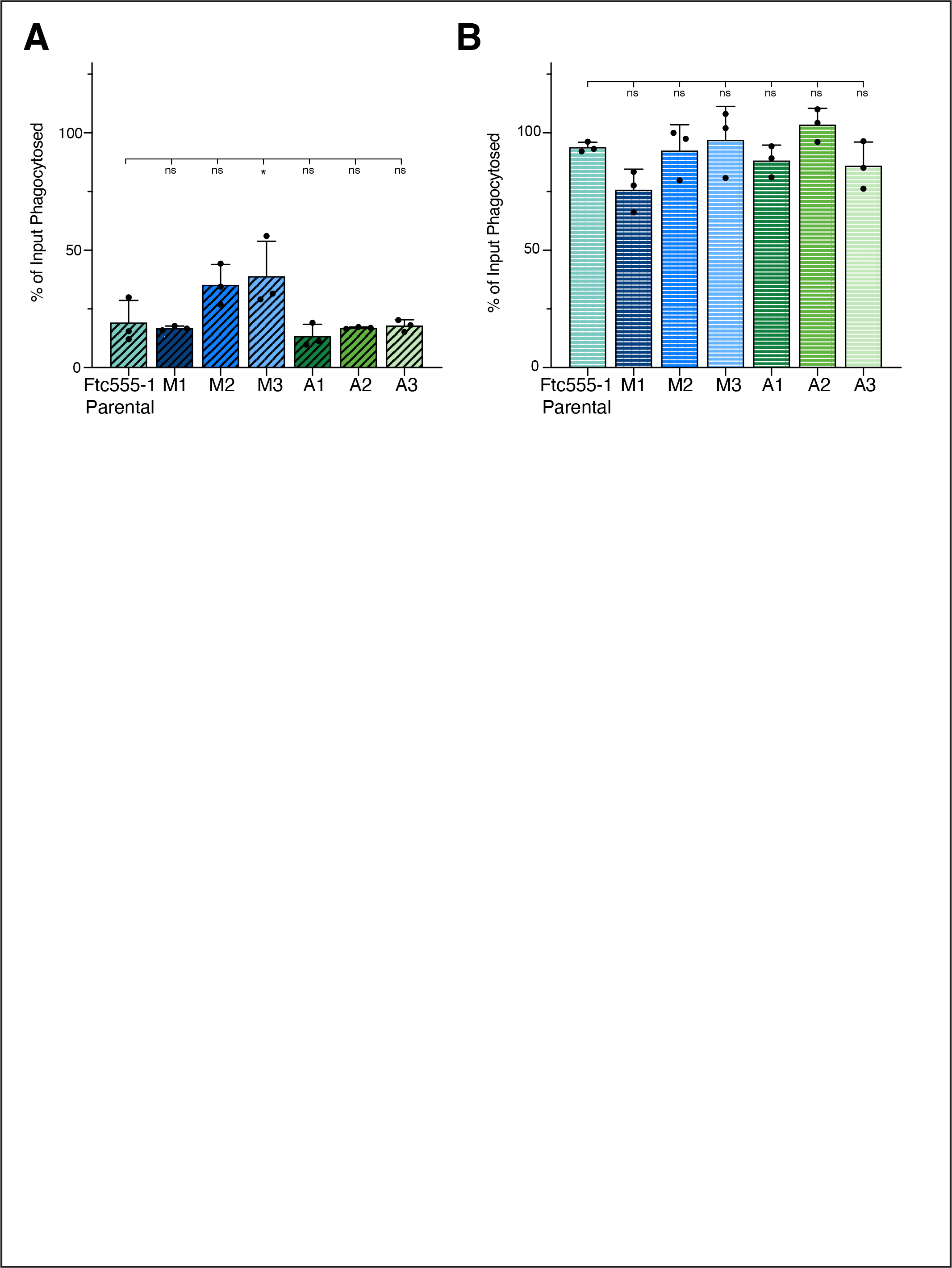
Differences in phagocytosis between evolved strains of Ftc555-1 do not affect replication phenotypes. The percent of cells from the input culture that were phagocytosed over one hour of incubation of *A. castellanii* cells (A) or mouse macrophages (B) calculated as above. Plotted data indicate the average values ± SD from one representative experiment. Each dot represents one replicate value from that experiment. Significance was assessed by comparison to the parental strain using ordinary one-way ANOVA followed by Dunnett’s multiple comparisons test. *p<0.05, ns, not significant.

**Supplemental Figure 3.**
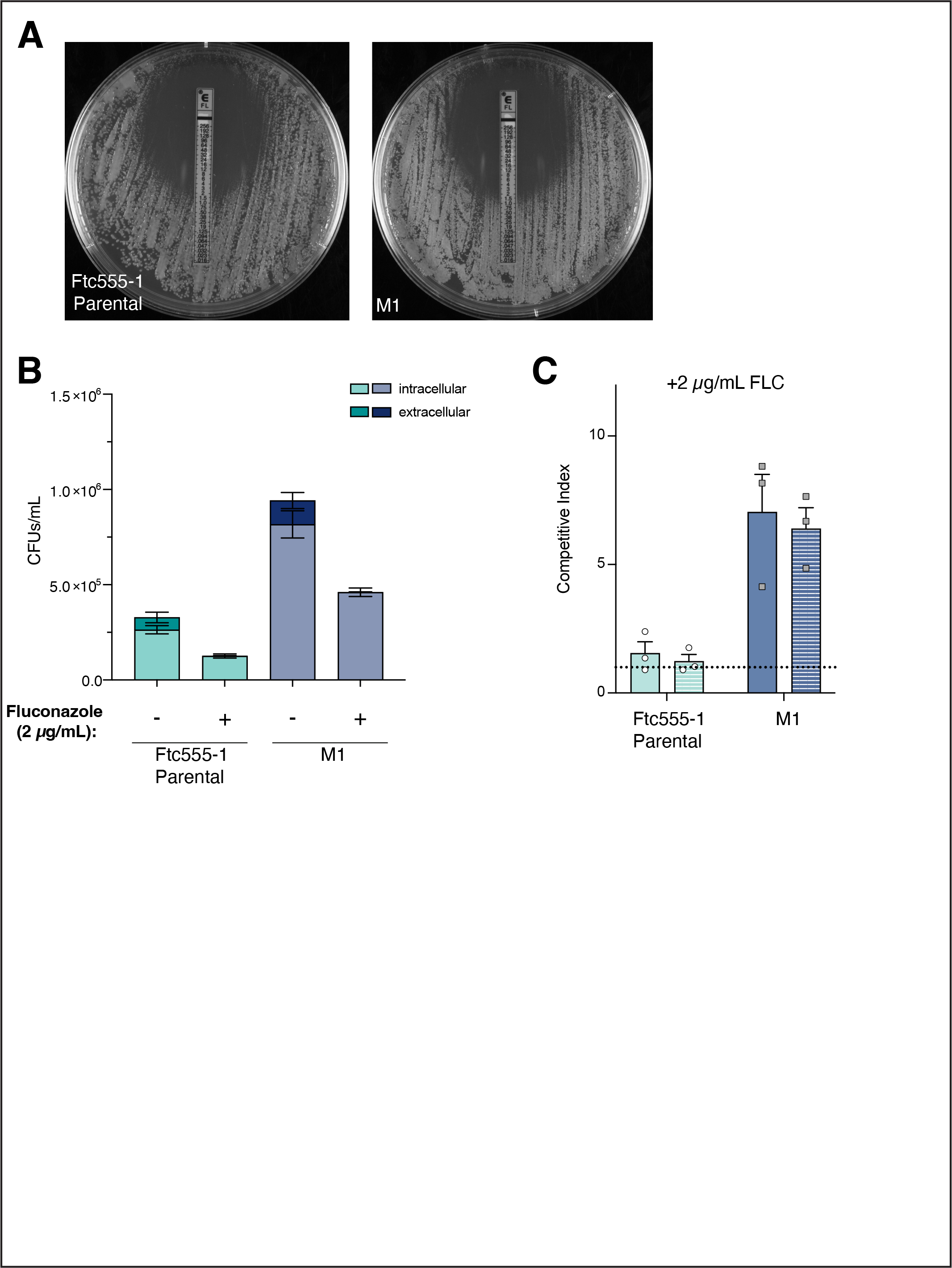
M1 has a fitness advantage in the macrophage intracellular niche. (A) Minimum inhibitory concentrations (MIC) for fluconazole were determined for the Ftc555-1 parental and M1 strains using an E-test. Both strains had identical MIC values by this method of 1.5-2 μg/mL. (B) The amount of intracellular (lighter shades) and extracellular (darker shades) growth in macrophage co-incubation experiments was determined with and without the addition of fluconazole to the culture media. Bars indicate average CFU counts ± SD from one representative experiment. Fluconazole eliminated most detectable extracellular growth for both Ftc555-1 and M1. Intracellular growth was also decreased in fluconazole treated wells (compare treated and untreated intracellular counts) but the increased replication of the M1 strain was preserved under both conditions. (C) Competitive indices for competition experiments performed with 2 μg/mL of fluconazole added to the media after phagocytosis. Competitive index was calculated as before. Bars indicate the average values ± SEM of three independent experiments, with dots indicating the average of three replicates from a single experiment. The dotted line shows a competitive index of one, which correlates with equal recovery of the NAT^R^ parental strain and the tested strain (i.e. no competitive advantage).

**Supplemental Figure 4.**
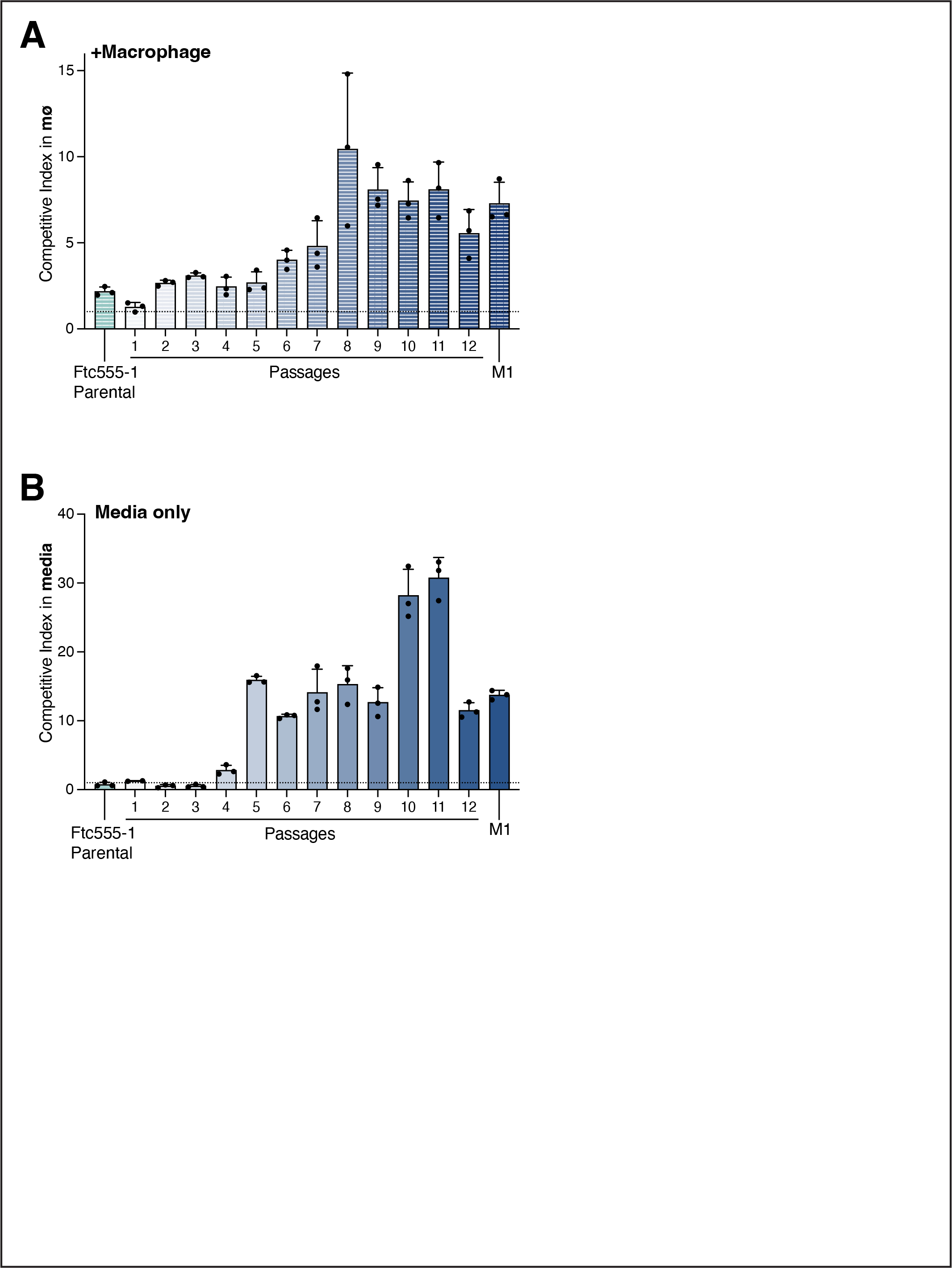
The M1 fitness advantage emerges following the acquisition of the *cac1-evo* allele. Competition experiments between populations of cells from each passage and the NAT^R^-labeled parental strain. Competitions were performed in macrophages (A) or in media-only (B). Competitive indices were calculated as before and the dotted line indicates a competitive index of one. Bars indicate the average of three replicate values from a single representive experiment.

## Supplemental Files.

**Supplemental File 1.** Strains used in this study.

**Supplemental File 2.** Oligos used in this study.

## References

1. Idnurm A, Bahn YS, Nielsen K, Lin X, Fraser JA, Heitman J. Deciphering the Model Pathogenic Fungus Cryptococcus Neoformans. Nat Rev Microbiol. 2005;3(10):753–64.

2. Montoya MC, Magwene PM, Perfect JR. Associations between Cryptococcus Genotypes, Phenotypes, and Clinical Parameters of Human Disease: A Review. J Fungi. 2021;7(4):260.

3. Rajasingham R, Govender NP, Jordan A, Loyse A, Shroufi A, Denning DW, et al. The global burden of HIV-associated cryptococcal infection in adults in 2020: a modelling analysis. Lancet Infect Dis. 2022;

4. Desjardins CA, Giamberardino C, Sykes SM, Yu CH, Tenor JL, Chen Y, et al. Population genomics and the evolution of virulence in the fungal pathogen Cryptococcus neoformans. Genome Res. 2017;27(7):1207–19.

5. Chen Y, Litvintseva AP, Frazzitta AE, Haverkamp MR, Wang L, Fang C, et al. Comparative analyses of clinical and environmental populations of Cryptococcus neoformans in Botswana. Mol Ecol. 2015;24(14):3559–71.

6. Litvintseva AP, Mitchell TG. Most Environmental Isolates of Cryptococcus neoformans var. grubii (Serotype A) Are Not Lethal for Mice. Infect Immun. 2009;77(8):3188–95.

7. Yu CH, Sephton-Clark P, Tenor JL, Toffaletti DL, Giamberardino C, Haverkamp M, et al. Gene Expression of Diverse Cryptococcus Isolates during Infection of the Human Central Nervous System. Mbio. 2021;12(6):e02313–21.

8. Yu CH, Chen Y, Desjardins CA, Tenor JL, Toffaletti DL, Giamberardino C, et al. Landscape of gene expression variation of natural isolates of Cryptococcus neoformans in response to biologically relevant stresses. Microb Genom. 2020;6(1).

9. Sephton-Clark P, Tenor JL, Toffaletti DL, Meyers N, Giamberardino C, Molloy SF, et al. Genomic variation across a clinical Cryptococcus population linked to disease outcome. Biorxiv. 2021;2021.11.22.469645.

10. Gerstein AC, Jackson KM, McDonald TR, Wang Y, Lueck BD, Bohjanen S, et al. Identification of Pathogen Genomic Differences That Impact Human Immune Response and Disease during Cryptococcus neoformans Infection. Mbio. 2019;10(4):e01440–19.

11. Casadevall A, Fu MS, Guimaraes A, Albuquerque P. The ‘Amoeboid Predator-Fungal Animal Virulence’ Hypothesis. J Fungi. 2019;5(1):10.

12. Casadevall A, Steenbergen JN, Nosanchuk JD. ‘Ready made’ virulence and ‘dual use’ virulence factors in pathogenic environmental fungi — the Cryptococcus neoformans paradigm. Curr Opin Microbiol. 2003;6(4):332–7.

13. Rayamajhee B, Willcox MDP, Henriquez FL, Petsoglou C, Subedi D, Carnt N. Acanthamoeba, an environmental phagocyte enhancing survival and transmission of human pathogens. Trends Parasitol. 2022;

14. Segal G, Shuman HA. Legionella pneumophila Utilizes the Same Genes To Multiply within Acanthamoeba castellanii and Human Macrophages. Infect Immun. 1999;67(5):2117–24.

15. Al-Quadan T, Price CT, Kwaik YA. Exploitation of evolutionarily conserved amoeba and mammalian processes by Legionella. Trends Microbiol. 2012;20(6):299–306.

16. Park JM, Ghosh S, O’Connor TJ. Combinatorial selection in amoebal hosts drives the evolution of the human pathogen Legionella pneumophila. Nat Microbiol. 2020;1–11.

17. O’Connor TJ, Adepoju Y, Boyd D, Isberg RR. Minimization of the Legionella pneumophila genome reveals chromosomal regions involved in host range expansion. Proc National Acad Sci. 2011;108(36):14733–40.

18. Steenbergen JN, Shuman HA, Casadevall A. Cryptococcus neoformans interactions with amoebae suggest an explanation for its virulence and intracellular pathogenic strategy in macrophages. Proc National Acad Sci. 2001;98(26):15245–50.

19. Chrisman CJ, Alvarez M, Casadevall A. Phagocytosis of Cryptococcus neoformans by, and Nonlytic Exocytosis from, Acanthamoeba castellanii⍰ †. Appl Environ Microb. 2010;76(18):6056–62.

20. Chrisman CJ, Albuquerque P, Guimaraes AJ, Nieves E, Casadevall A. Phospholipids Trigger Cryptococcus neoformans Capsular Enlargement during Interactions with Amoebae and Macrophages. Plos Pathog. 2011;7(5):e1002047.

21. Derengowski L da S, Paes HC, Albuquerque P, Tavares AHFP, Fernandes L, Silva-Pereira I, et al. The Transcriptional Response of Cryptococcus neoformans to Ingestion by Acanthamoeba castellanii and Macrophages Provides Insights into the Evolutionary Adaptation to the Mammalian Host. Eukaryot Cell. 2013;12(5):761–74.

22. Rhodes J, Beale MA, Vanhove M, Jarvis JN, Kannambath S, Simpson JA, et al. A Population Genomics Approach to Assessing the Genetic Basis of Within-Host Microevolution Underlying Recurrent Cryptococcal Meningitis Infection. G3 Genes Genomes Genetics. 2017;7(4):1165–76.

23. Ormerod KL, Morrow CA, Chow EWL, Lee IR, Arras SDM, Schirra HJ, et al. Comparative Genomics of Serial Isolates of Cryptococcus neoformans Reveals Gene Associated With Carbon Utilization and Virulence. G3 Genes Genomes Genetics. 2013;3(4):675–86.

24. Chen Y, Farrer RA, Giamberardino C, Sakthikumar S, Jones A, Yang T, et al. Microevolution of Serial Clinical Isolates of Cryptococcus neoformans var. grubii and C. gattii. Mbio. 2017;8(2):e00166–17.

25. Brunke S, Seider K, Fischer D, Jacobsen ID, Kasper L, Jablonowski N, et al. One Small Step for a Yeast - Microevolution within Macrophages Renders Candida glabrata Hypervirulent Due to a Single Point Mutation. Plos Pathog. 2014;10(10):e1004478.

26. Wartenberg A, Linde J, Martin R, Schreiner M, Horn F, Jacobsen ID, et al. Microevolution of Candida albicans in Macrophages Restores Filamentation in a Nonfilamentous Mutant. Plos Genet. 2014;10(12):e1004824.

27. Hu G, Chen SH, Qiu J, Bennett JE, Myers TG, Williamson PR. Microevolution During Serial Mouse Passage Demonstrates FRE3 as a Virulence Adaptation Gene in Cryptococcus neoformans. Mbio. 2014;5(2):e00941–14.

28. Sephton-Clark P, McConnell SA, Grossman N, Baker R, Dragotakes Q, Fan Y, et al. Human and murine Cryptococcus neoformans infection selects for common genomic changes in an environmental isolate. Biorxiv. 2022;2022.04.12.487930.

29. Alvarez M, Casadevall A. Phagosome Extrusion and Host-Cell Survival after Cryptococcus neoformans Phagocytosis by Macrophages. Curr Biol. 2006;16(21):2161–5.

30. Ma H, Croudace JE, Lammas DA, May RC. Expulsion of Live Pathogenic Yeast by Macrophages. Curr Biol. 2006;16(21):2156–60.

31. Seoane PI, May RC. Vomocytosis: What we know so far. Cell Microbiol. 2020;22(2):e13145.

32. Alspaugh JA, Pukkila-Worley R, Harashima T, Cavallo LM, Funnell D, Cox GM, et al. Adenylyl Cyclase Functions Downstream of the Gα Protein Gpa1 and Controls Mating and Pathogenicity of Cryptococcus neoformans. Eukaryot Cell. 2002;1(1):75–84.

33. Hicks JK, D’Souza CA, Cox GM, Heitman J. Cyclic AMP-Dependent Protein Kinase Catalytic Subunits Have Divergent Roles in Virulence Factor Production in Two Varieties of the Fungal Pathogen Cryptococcus neoformans. Eukaryot Cell. 2004;3(1):14–26.

34. D’Souza CA, Alspaugh JA, Yue C, Harashima T, Cox GM, Perfect JR, et al. Cyclic AMP-Dependent Protein Kinase Controls Virulence of the Fungal Pathogen Cryptococcus neoformans. Mol Cell Biol. 2001;21(9):3179–91.

35. Alspaugh JA, Perfect JR, Heitman J. Cryptococcus neoformans mating and virulence are regulated by the G-protein α subunit GPA1 and□cAMP. Gene Dev. 1997;11(23):3206–17.

36. Dambuza IM, Drake T, Chapuis A, Zhou X, Correia J, Taylor-Smith L, et al. The Cryptococcus neoformans Titan cell is an inducible and regulated morphotype underlying pathogenesis. Plos Pathog. 2018;14(5):e1006978.

37. Caza M, Kronstad JW. The cAMP/Protein Kinase A Pathway Regulates Virulence and Adaptation to Host Conditions in Cryptococcus neoformans. Front Cell Infect Mi. 2019;9:212.

38. Bahn YS, Hicks JK, Giles SS, Cox GM, Heitman J. Adenylyl Cyclase-Associated Protein Aca1 Regulates Virulence and Differentiation of Cryptococcus neoformans via the Cyclic AMP-Protein Kinase A Cascade. Eukaryot Cell. 2004;3(6):1476–91.

39. Casadevall A, Rosas AL, Nosanchuk JD. Melanin and virulence in Cryptococcus neoformans. Curr Opin Microbiol. 2000;3(4):354–8.

40. Zaragoza O, Casadevall A. Experimental modulation of capsule size in Cryptococcus neoformans. Biol Proced Online. 2004;6(1):10–5.

41. Zaragoza O, Fries BC, Casadevall A. Induction of Capsule Growth in Cryptococcus neoformans by Mammalian Serum and CO 2. Infect Immun. 2003;71(11):6155–64.

42. Granger DL, Perfect JR, Durack DT. Virulence of Cryptococcus neoformans. Regulation of capsule synthesis by carbon dioxide. J Clin Invest. 1985;76(2):508–16.

43. Mukaremera L, McDonald TR, Nielsen JN, Molenaar CJ, Akampurira A, Schutz C, et al. The Mouse Inhalation Model of Cryptococcus neoformans Infection Recapitulates Strain Virulence in Humans and Shows that Closely Related Strains Can Possess Differential Virulence. Infect Immun. 2019;87(5).

44. Hommel B, Mukaremera L, Cordero RJB, Coelho C, Desjardins CA, Sturny-Leclère A, et al. Titan cells formation in Cryptococcus neoformans is finely tuned by environmental conditions and modulated by positive and negative genetic regulators. Plos Pathog. 2018;14(5):e1006982.

45. Slev PR, Potts WK. Disease consequences of pathogen adaptation. Curr Opin Immunol. 2002;14(5):609–14.

46. Didelot X, Walker AS, Peto TE, Crook DW, Wilson DJ. Within-host evolution of bacterial pathogens. Nat Rev Microbiol. 2016;14(3):150–62.

47. Duffy S, Shackelton LA, Holmes EC. Rates of evolutionary change in viruses: patterns and determinants. Nat Rev Genet. 2008;9(4):267–76.

48. Lenski RE. Experimental evolution and the dynamics of adaptation and genome evolution in microbial populations. Isme J. 2017;11(10):2181–94.

49. McClelland EE, Adler FR, Granger DL, Potts WK. Major histocompatibility complex controls the trajectory but not host-specific adaptation during virulence evolution of the pathogenic fungus Cryptococcus neoformans. Proc Royal Soc Lond Ser B Biological Sci. 2004;271(1548):1557–64.

50. Fu MS, Liporagi-Lopes LC, Santos SR dos, Tenor JL, Perfect JR, Cuomo CA, et al. Amoeba Predation of Cryptococcus neoformans Results in Pleiotropic Changes to Traits Associated with Virulence. Mbio [Internet]. 2021;12(2):e00567–21. Available from: https://journals.asm.org/doi/epdf/10.1128/mBio.00567-21

51. Fan Y, Lin X. Multiple Applications of a Transient CRISPR-Cas9 Coupled With Electroporation (TRACE) System in the Cryptococcus neoformans Species Complex. Genetics. 2018;208(4):genetics.300656.2018.

52. Huang MY, Joshi MB, Boucher MJ, Lee S, Loza LC, Gaylord EA, et al. Short homology-directed repair using optimized Cas9 in the pathogen Cryptococcus neoformans enables rapid gene deletion and tagging. Genetics. 2021;220(1).

53. Upadhya R, Lam WC, Maybruck BT, Donlin MJ, Chang AL, Kayode S, et al. A fluorogenic C. neoformans reporter strain with a robust expression of m-cherry expressed from a safe haven site in the genome. Fungal Genet Biol. 2017;108:13–25.

54. Drenkard E, Richter BG, Rozen S, Stutius LM, Angell NA, Mindrinos M, et al. A Simple Procedure for the Analysis of Single Nucleotide Polymorphisms Facilitates Map-Based Cloning in Arabidopsis. Plant Physiol. 2000;124(4):1483–92.

55. Koren S, Walenz BP, Berlin K, Miller JR, Bergman NH, Phillippy AM. Canu: scalable and accurate long-read assembly via adaptive k-mer weighting and repeat separation. Genome Res. 2017;27(5):722–36.

56. Walker BJ, Abeel T, Shea T, Priest M, Abouelliel A, Sakthikumar S, et al. Pilon: An Integrated Tool for Comprehensive Microbial Variant Detection and Genome Assembly Improvement. Plos One. 2014;9(11):e112963.

57. Jin JJ, Yu WB, Yang JB, Song Y, dePamphilis CW, Yi TS, et al. GetOrganelle: a fast and versatile toolkit for accurate de novo assembly of organelle genomes. Genome Biol. 2020;21(1):241.

58. Shumate A, Salzberg SL. Liftoff: accurate mapping of gene annotations. Bioinformatics. 2021;37(12):btaa1016.

59. Md V, Misra S, Li H, Aluru S. Efficient Architecture-Aware Acceleration of BWA-MEM for Multicore Systems. 2019 Ieee Int Parallel Distributed Process Symposium Ipdps. 2019;00:314–24.

60. Bolger AM, Lohse M, Usadel B. Trimmomatic: a flexible trimmer for Illumina sequence data. Bioinformatics. 2014;30(15):2114–20.

61. Danecek P, Bonfield JK, Liddle J, Marshall J, Ohan V, Pollard MO, et al. Twelve years of SAMtools and BCFtools. Gigascience. 2021;10(2):giab008.

62. McKenna A, Hanna M, Banks E, Sivachenko A, Cibulskis K, Kernytsky A, et al. The Genome Analysis Toolkit: A MapReduce framework for analyzing next-generation DNA sequencing data. Genome Res. 2010;20(9):1297–303.

63. Cingolani P, Platts A, Wang LL, Coon M, Nguyen T, Wang L, et al. A program for annotating and predicting the effects of single nucleotide polymorphisms, SnpEff. Fly. 2012;6(2):80–92.

64. Thorvaldsdóttir H, Robinson JT, Mesirov JP. Integrative Genomics Viewer (IGV): high-performance genomics data visualization and exploration. Brief Bioinform. 2013;14(2):178–92.

65. Nichols CB. Visualization and Documentation of Capsule and Melanin Production in Cryptococcus neoformans. Curr Protoc. 2021;1(1):e27.

66. Schindelin J, Arganda-Carreras I, Frise E, Kaynig V, Longair M, Pietzsch T, et al. Fiji: an open-source platform for biological-image analysis. Nat Methods. 2012;9(7):676–82.

67. Steenwyk JL, Rokas A. Treehouse: a user-friendly application to obtain subtrees from large phylogenies. Bmc Res Notes. 2019;12(1):541.

